# Molecular basis for recognition of *Listeria* cell wall teichoic acid by the pseudo-symmetric SH3b-like repeats of a bacteriophage endolysin

**DOI:** 10.1101/2020.06.05.136911

**Authors:** Yang Shen, Ioanna Kalograiaki, Alessio Prunotto, Matthew Dunne, Samy Boulos, Nicholas M.I. Taylor, Eric Sumrall, Marcel R. Eugster, Rebecca Martin, Alicia Julian-Rodero, Benjamin Gerber, Petr G. Leiman, Margarita Menéndez, Matteo Dal Peraro, Francisco Javier Cañada, Martin J. Loessner

## Abstract

Endolysins are bacteriophage-encoded peptidoglycan hydrolases targeting the cell wall of host bacteria via their cell wall-binding domains (CBDs). The molecular basis for selective recognition of surface carbohydrate ligands by CBDs remains elusive. Here, we describe, in atomic detail, the interaction between the *Listeria* phage endolysin domain CBD500 and its cell wall teichoic acid (WTA) ligands. We show that 3’ *O*-acetylated GlcNAc residues integrated into the WTA polymer chain are the key epitope recognized by a CBD binding cavity located at the interface of tandem copies of *beta*-barrel, pseudo-symmetric SH3b-like repeats. This cavity consists of multiple aromatic residues making extensive interactions with two GlcNAc acetyl groups via hydrogen bonds and van der Waals contacts, while permitting the docking of the diastereomorphic ligands. The multidisciplinary approach described here delineates a previously unknown recognition mechanism by which a phage endolysin specifically recognizes and targets WTA, suggesting an adaptable model for regulation of endolysin specificity.

## Introduction

Bacteriophage-encoded endolysins are peptidoglycan hydrolases that digest the bacterial cell wall and induce bacterial lysis at the final stage of the phage lytic cycle^1^. Although they naturally work from within the infected cells, exogenous application of recombinant enzyme to susceptible bacteria can likewise lead to rapid degradation of their cell wall, termed lysis from-without^2, 3^. Due to their exceptional specificity and efficacy, phage endolysins are increasingly considered as alternatives to antibiotics for combating multidrug-resistant bacteria^4–6^.

The vast majority of endolysins from bacteriophages infecting Gram-positive bacteria possess modular architectures, encompassed in at least one enzymatically-active domain (EAD) and one cell wall-binding domain (CBD)^7^. The *N*-terminal EAD is responsible for cleaving a particular peptidoglycan (PG) bond, and the *C*-terminal CBD usually holds an independent regulatory function and determines the lytic spectrum upon targeting specific cell-wall elements distributed in a genus-, species- or strain-specific manner. The binding ligands for CBDs are usually of peptidoglycan or carbohydrate nature^8–10^. Given their binding specificity, endolysins and in particular the CBDs constitute highly attractive scaffolds for antibacterial protein engineering^11, 12^.

*Listeria* phage A500 encodes the endolysin Ply500, which comprises L-alanoyl-D-glutamate endopeptidase activity against *Listeria spp.* serovars 4, 5, and 6^10^. Notably, Ply500 is able to rapidly lyse all serovar 4b strains, which are responsible for the vast majority of listeriosis outbreaks^13, 14^, rendering Ply500 an appealing anti-*Listeria* agent to use in food safety applications^15–17^. Ply500 displays a modular architecture comprising largely independent CBD and EAD moieties connected *via* a short and flexible linker peptide (**Fig. 1a**)^18^. Although both modules seem to be required for full activity of this endolysin on its natural murein substrate^10, 18^, the isolated *C*-terminal domain (termed CBD500) binds live bacteria cells^10, 19^, as well as to purified wall teichoic acid (WTA)^20^ with high affinity.

**Figure 1.**
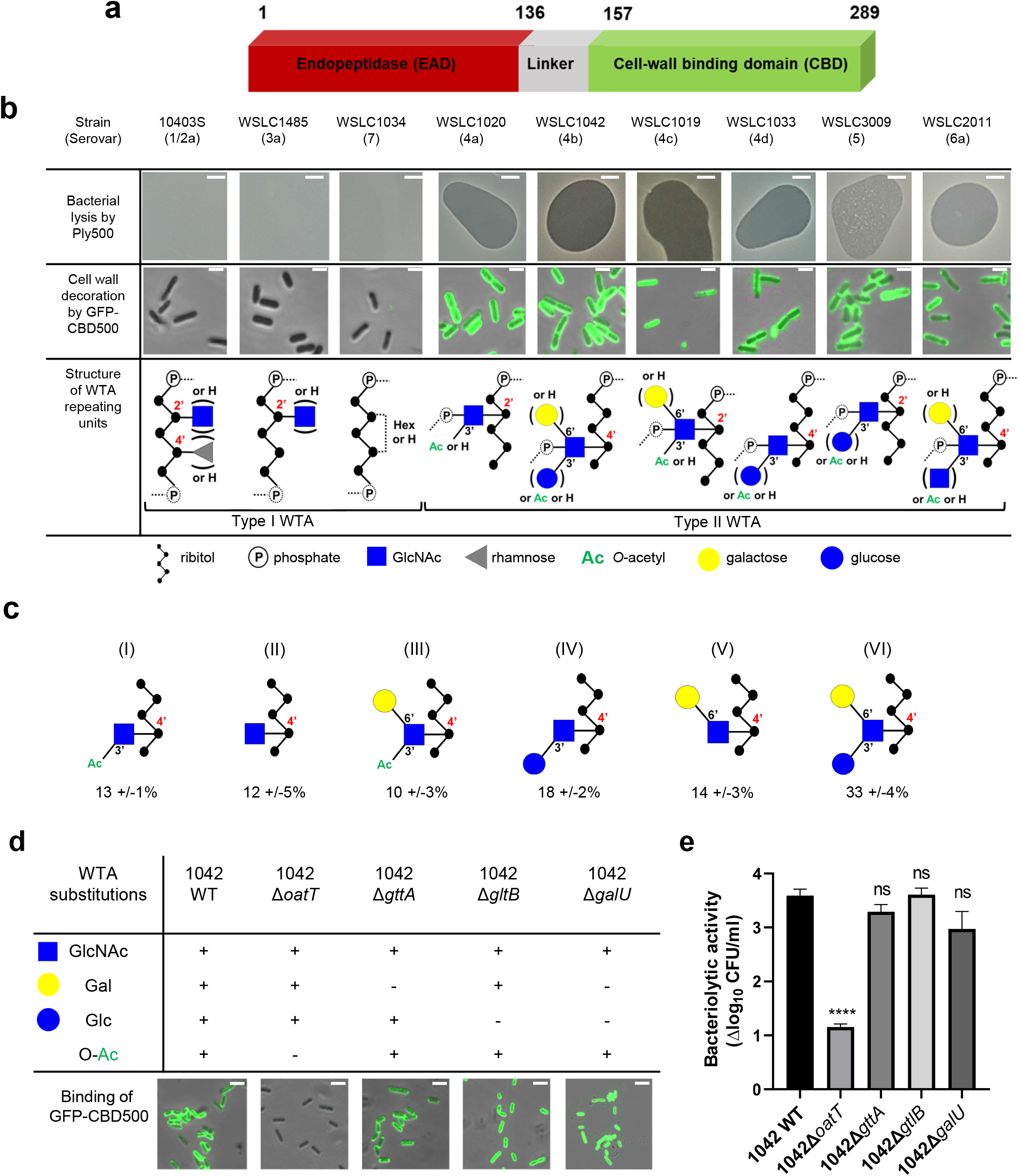
Correlation of *Listeria* WTA structure with the bacteriolytic activity of Ply500 endolysin and the binding activity of its C-terminal domain CBD500. **a**, Modular architecture of the full-length Ply500. **b**, Lytic activity of full-length Ply500 detected by spot-on-lawn assay (upper panel; scale bar 300 μm), and binding specificity of green-fluorescent-protein labeled CBD500 (GFP-CBD500) against *Listeria* strains (middle panel; scale bar 2 μm) decorated with distinct types of WTA^20^. Images are representative of three experiments, and contrast was adjusted for clarity. **c**, Relative abundance (%) of the repeating unit variants within *Listeria* WTAs derived from strain WSLC1042 (WTA_1042_), determined by UPLC-MS analysis. Latin numerals (I-VI) denote the repeating unit species of WTA_1042_ identified. Data represent means ± SEM; n = 3. **d**, Binding of GFP-CBD500 to wild-type WSLC1042 and mutants lacking specific substituents in the backbone of the WTA repeating unit. Scale bar represents 1 μm. Images are representative of three experiments, and contrast was adjusted for clarity. **e**, Reduction of bacterial colony counts (Δ log_10_CFU ml^−1^) after 1 h incubation of WSLC1042 and selected WTA mutants in the presence of 100 μg/ml of Ply500. Results are the mean ± SEM of three experiments. Statistical significance was assessed using one-way ANOVA test (****p<0.0001; ns: not significant).

In *Listeria*, WTA polymers accounts for up to 60% of the dry weight of the cell-wall, and are covalently bound to *N*-acetylmuramic acid (MurNAc) of the PG by a phosphodiester bond through a conserved linkage unit^21^. The WTA chains extend beyond the PG layer, comprising 20 to 40 repeating units that are highly variable due to further glycosidic substitutions or modifications^22^. Depending on their backbone units, two types of WTA exist (**Fig. 1b**): Type I WTAs possess poly-[5)-Rbo-1-P-(O→]_*n*_ chains of ribitol phosphate (Rbo-P) units, while type II WTAs feature repeating units of GlcNAc-Rbo-P, following the pattern [4)-GlcNAc-(*β*1→2/4)-Rbo-1-P-(O→]_*n*_. WTAs represent the *O*-antigen determinants in *Listeria spp.,* and confer serovar diversity along with some *H*-antigens^23^. We previously demonstrated that fluorescently labelled CBDs with distinct recognition and binding pattern could differentiate *Listeria* cells down to the serovar level^24, 25^. In particular, CBD500 was shown to specifically decorate *Listeria* strains carrying type II WTAs equipped with 3’ *O*-acetylated GlcNAc, while tolerating further hexose substitution of the integrated GlcNAc moiety (**Fig. 1b**)^20^. Yet, the underlying recognition mechanism remains elusive with respect to polymer length, GlcNAc/ribitol connectivity, inherent flexibility and microheterogeneity (due to non-uniform, variable decoration of the repeating units). Thus, the attainment of an atomic-scale model required the combination of multiplex ligand- and receptor-based methods.

We used a state-of-the-art multidisciplinary approach to elucidate the mechanism underpinning the recognition of *Listeria* WTAs by CBD500 from both the ligand and protein perspectives. Employing X-ray crystallography, we solved the structure of CBD500 to 1.6 Å resolution, which comprises two copies of *beta*-barrel SH3b-like repeats held together by means of swapped *beta*-strands. Saturation Transfer Difference (STD) NMR^26^ studies of CBD500 in complex with full length WTAs derived from a selected panel of *Listeria* strains and mutants, highlighted the prerequisite of GlcNAc 3’ *O*-acetylation for interaction with CBD500. “Blind” molecular docking on the solved structure suggested three putative binding pockets whose involvement in binding was furtherly assessed by site-directed mutagenesis and functionality assays. STD NMR-driven molecular dynamics simulations and isothermal titration calorimetry (ITC) aided to model the interaction, which is driven by hydrogen bond networks and van der Waals contacts. For the first time, we provide the atomic evidence that swapped SH3b-like repeats found in a *Listeria* phage endolysin permit the selective recognition of carbohydrate ligands^10, 27^ instead of PG-related peptides^28–30^. This pioneering study fosters the existing knowledge on WTA recognition by endolysins, and provides evidence on the diversification of the SH3b domain folding and binding site. Altogether, our findings pave the way towards developing novel diagnostic and therapeutic strategies targeting pathogenic bacteria.

## Results

### Ply500 targets *Listeria* WTAs by recognizing *O*-acetylated GlcNAc residues

Spot-on-lawn assays and fluorescence decoration assays confirmed previous findings that Ply500 specifically targets bacterial cell walls displaying type II WTAs (**Fig. 1b**), all of which feature *O*-acetylated GlcNAc integrated in the polymeric ribitol chain^10, 20^. To dissect the role of each respective modification of WTA in CBD500 binding, a series of WTA gene deletion mutants^3^ derived from the *L. monocytogenes* serovar 4b strain WSLC 1042 were employed. In the WT strain, WTA_1042_ features an unequal ratio of six monomer variants, determined by UPLC-MS analysis (**Fig. 1c**). Knock-out of *oatT* resulted in loss of WTA *O*-acetylation (**Supplementary Fig. 1a**) and CBD500 binding, whereas deletion of *gttA* (led to a loss of Gal decoration) did not change the binding (**Fig. 1d**). Interestingly, loss of Glc in both *galU* and *gltB* knock-outs led to slightly enhanced binding, possibly owing to the higher degree of *O*-acetylation found in their WTAs (**Supplementary Fig. 1b**). Consistent with the loss of CBD500 binding, 1042Δ*oatT* mutant cells were found to be significantly less sensitive to Ply500-mediated lysis as opposed to other WTA mutants (**Fig. 1e**).

Next, we sought to validate bacterial binding using CBD500 alone and extracted WTA polymers. As such, we immobilized CBD500 onto a surface plasmon resonance (SPR) chip, and used wild type and mutant WTA as the analytes. The SPR analysis revealed that the association of WTA with CBD500 is abolished for WTA lacking the *O*-acetylation (**Supplementary Fig. 1c**). Likewise, binding of CBD500 to *Listeria ivanovii* WSLC 3009 (SV 5) was also prevented upon loss of GlcNAc 3’ *O*-acetylation (**Supplementary Fig. 1d,e**). In summary, these results indicate that CBD500 specifically recognizes *O*-acetylated WTA as its ligand.

### Identification of the carbohydrate epitope recognized by CBD500 with STD-NMR

Following the assignment of the NMR signals of WTAs derived from WT strains and mutants (**Supplementary Fig. S2a-c**, **Table 1** **and 2**) and estimation of their mean molecular size by diffusion ordered spectroscopy^32^ (DOSY) (14.7 and 17.7 kDa for WTA_1042_ and WTA_1020_) respectively, (**Supplementary Fig. S2d, e**), their recognition by CBD500 was investigated by STD-NMR. This technique is based on selective irradiation of protein protons and subsequent detection of magnetization transfer to the ligand, allowing for uncovering of middle-range affinity binding, ligand screening, and epitope mapping^33^. Measurements carried out in the presence of CBD500 gave rise to STD-positive peaks for both WTA_1042_ and WTA_1020_ (**Fig. 2a** and **Supplementary Table 3**), confirming specific binding of CBD500 to these structures. Resonances with chemical shifts (δ) in the range 2-2.13 ppm were unambiguously assigned to the methyl protons of GlcNAc units. For both WTA_1042_ and WTA_1020_, the strongest STD signals (*ca.* 16-25%) compared to the off-resonance spectra were observed for the *N*-Ac- and *O*-Ac-methyl protons of the 3’OAc-GlcNAc, demonstrating that this unit is the primary docking point for the protein (**Fig. 2a**). Although several proton resonances of the 3’OAc-GlcNAc moiety are located in the crowded spectral region between 3.5-4.3 ppm, where partial resonance overlap complicates the assignment (**Supplementary Table 3**), peaks belonging to H3’ of the 3’ O-acetylated unit (~5.2 ppm) were clearly distinguished in the STD difference spectrum, claiming for a strong engagement of the GlcNAc ring in binding. For WTA_1042_, and to a lesser extent for WTA_1020_, high magnetization transfer was observed for the 3’OAc-GlcNAc H4’ protons (**Fig. 2a** and **Supplementary Table 3**) located on the opposite face of the pyranose ring. In striking contrast, the *α*Gal and Rbo protons solely received medium to low saturation transfer, suggesting larger distances to the protein surface (**Fig. 2b**), again pointing to the GlcNAc moiety as the key interaction residue. Only negligible transfer was observed to the *N*-Ac-methyl protons of GlcNAc lacking 3’*O*-acetylation (less than 2% absolute values for WTA1042) discarding a significant involvement of these units in complex formation.

**Table 1.** *Listeria* cell wall binding capacity of GFP-CBD500 alanine mutants and ITC binding-thermodynamics to WTA_1042._

| Constructs | Binding to *Listeria*<sup>a</sup> | N<sup>b</sup> | K<sub>b</sub><br>(M<sup>-1</sup>) | -ΔH<br>(kcal/mol) | -ΔG<br>(kcal/mol) | TΔS<br>(kcal/mol) |
| --- | --- | --- | --- | --- | --- | --- |
| WT | + | 1.88±0.03 | (3.5±0.2)<br>× 10<sup>4</sup> | 9.5±0.1 | 6.20±0.03 | -3.3±0.1 |
| K162A | + | 2.1±0.2 | (2.8±0.4)<br>× 10<sup>4</sup> | 6.7 ±0.2 | 6.07±0.08 | -0.6±0.3 |
| Y197A | (+) | 1.8±0.1 | (3.3±0.4)<br>× 10<sup>4</sup> | 4.3 ±0.2 | 6.16±0.07 | 1.9±0.3 |
| Y211A | (+) | 2.2±0.2 | (2.5±0.2)<br>× 10<sup>4</sup> | 3.0 ± 0.2 | 6.00±0.05 | 3.0±0.25 |
| K246A | + | 2.0±0.1 | (2.3±0.2)<br>× 10<sup>4</sup> | 10.5 ±0.2 | 5.95±0.05 | -<br>4.55±0.45 |
| S249A | (+) | 1.5±0.1 | (3.6±0.5)<br>× 10<sup>4</sup> | 8.2±0.2 | 6.30±0.03 | -1.9±0.2 |
| K251D | + | 1.0±0.1 | (3.5±0.5)<br>× 10<sup>4</sup> | 8±1 | 6.20±0.08 | -1.8±1 |
| K289A | + | 2.0±0.2 | (4.8±0.7)<br>× 10<sup>4</sup> | 5.7±0.2 | 6.39±0.09 | 0.7±0.3 |
| K246A_N248A | + | 2.3 ±0.5 | (2.9±0.3)<br>× 10<sup>4</sup> | 8.4±0.3 | 6.09±0.06 | -2.3±0.4 |
| I252A | (+) | Protein aggregation prevented quantification of ITC data |  |  |  |  |
| W254A | - | Extremely weak affinity not quantifiable by ITC |  |  |  |  |
| F175A<sup>c</sup> | - | No binding detected by ITC |  |  |  |  |
| W198A | - | Extremely weak affinity not quantifiable by ITC |  |  |  |  |
| W242A<sup>c</sup> | - | No binding detected by ITC |  |  |  |  |
| N248A<sup>d</sup> | - | Extremely weak affinity not quantifiable by ITC |  |  |  |  |
| GFP | - | No binding detected by ITC |  |  |  |  |

<sup>a</sup>Binding was visualized by fluorescence microscopy using the GFP-CBD500 construct and its variants: '+', strong binding; '(+)', weak binding; '-', no binding.
<sup>b</sup>N is the number of CBD molecules bound per WTA chain, K<sub>b</sub>, is the binding constant per monomer of CBD bound to WTA as measured by ITC, and ΔG (-RT lnK<sub>b</sub>), ΔH and ΔS represent the corresponding thermodynamic parameters.
<sup>c</sup>ITC measured with CBD500 mutants. <sup>d</sup>ITC measured with CBD500 and GFP-CBD500 mutants.

**Figure 2.**
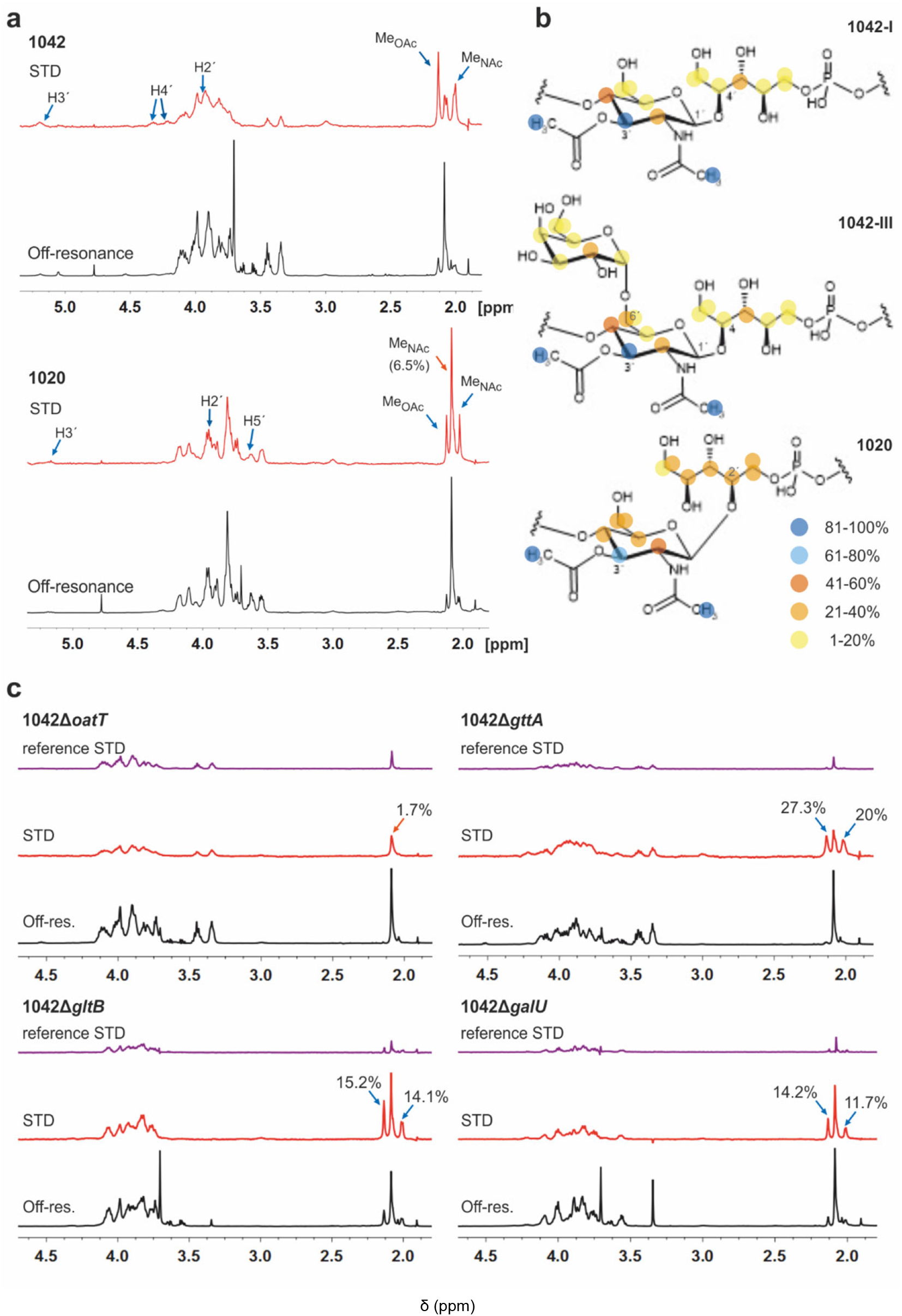
WTA_1042_ and WTA_1020_ epitopes recognized by CBD500. **a**, Representative Saturation Transfer Difference (STD) NMR (red, scaled 16x) and off-resonance (black) spectra of WTA_1042_ and WTA_1020_ recorded in presence of CBD500 upon irradiation at 7.1 and 100 ppm, respectively (1:25 CBD:WTA mixtures containing 30 μΜ CBD500 and approx. 0.75 mM WTA repeating units). Select STD signals are highlighted. Blue arrows point to *O*-acetylated GlcNAc protons and orange arrows to GlcNAc protons, bearing –OH or –OGlc at C3’. **b,** STD epitopes for WTA_1042-I_, WTA_1042-III_ and WTA_1020_ *O*-acetylated GlcNAc moieties as derived from at least three independent experiments; blue and yellow spheres indicate higher and lower saturation transfer intensities. **c**, Representative STD (red, scaled 16x) and off-resonance (black) spectra of CBD500-WTA_1042_ complexes where WTA were derived from *Lm* 1042 mutants Δ*oatT* Δ*gttA*, Δ*gltB* and Δ*galU* as indicated; reference STD spectra acquired in absence of CBD500 are also depicted for all mutants in purple (ligand-only control, same scale as STD in presence of protein). Select STD signals are highlighted and absolute STD values are given for the methyl-group protons of relevance. All spectra were acquired as above and the chemical shifts are measured in ppm. Detailed information on STD absolute and normalized values is summarized in Supplementary **Tables 3** and **4**.

To establish the minimum WTA epitope required for CBD500-WTA interaction and investigate the relevant contributions of 3’ *O*-acetylation or hexose (3’ -OGlc or 6’ -OGal) substitutions of the GlcNAc moiety, STD-NMR experiments were also performed using WTA isolated from defined WTA mutants (**Fig. 1d**). Lack of 3’ *O*-acetylation completely abolished recognition by CBD500 (1042Δ*oatT*, **Fig. 2c**), in line with the previous observations (**Supplementary Fig. 1c**). Indeed, the absolute STD value measured for the GlcNAc *N*-Ac-methyl protons was equivalent to the non-specific ligand saturation observed in absence of CBD500 (reference STD, **Fig. 2c**). For all three variant WTAs derived from the 1042Δ*gttA*, Δ*gltB* or Δ*galU* mutants, magnetization transfer values were of similar intensity (**Supplementary Table 4**), suggesting that 3’O-Glc and 6’O-Gal modifications are not essential for CBD500 recognition. Overall, findings obtained by NMR established 3’OAc-GlcNAc moiety as the key epitope for CBD500, and, in the case of WTA_1042_, indicate that Gal substitution at position 6’ is well tolerated.

### Crystal structure of CBD500

To further our understanding of the molecular interactions between CBD500 and WTA, we determined the crystal structure of His-CBD500 (aa 133-289) at 1.6 Å resolution by molecular replacement using the CBD of PlyPSA (PDB ID: 1XOV) as a search model (structure solution statistics provided in **Supplementary table 5**). As described for Ply PSA^27^, CBD500 consists of two adjoined SH3b-like (pfam PF08239) domains connected through structurally swapped β-sheets (**Fig. 3a**). The first β-strand (β1p/β1’d) forms a part of the proximal domain before connecting to the distal domain (β2d to β8d/β1’p) with the end of β8d connected to β2p of the proximal domain, which concludes with β6p. Only the distal domain features the α-helix turn (α1) and an additional β-strand (β7) as found in other SH3b domains^34, 35^. The two RT loops^36^ (connecting β2p-β3p and β2d-β3d) that are distinctive features of SH3 and SH3b repeats are positioned facing each other at the interface between the domains. For a simplified view, a topological diagram of the CBD500 secondary structure is shown in **Fig. 3b**.

**Figure 3.**
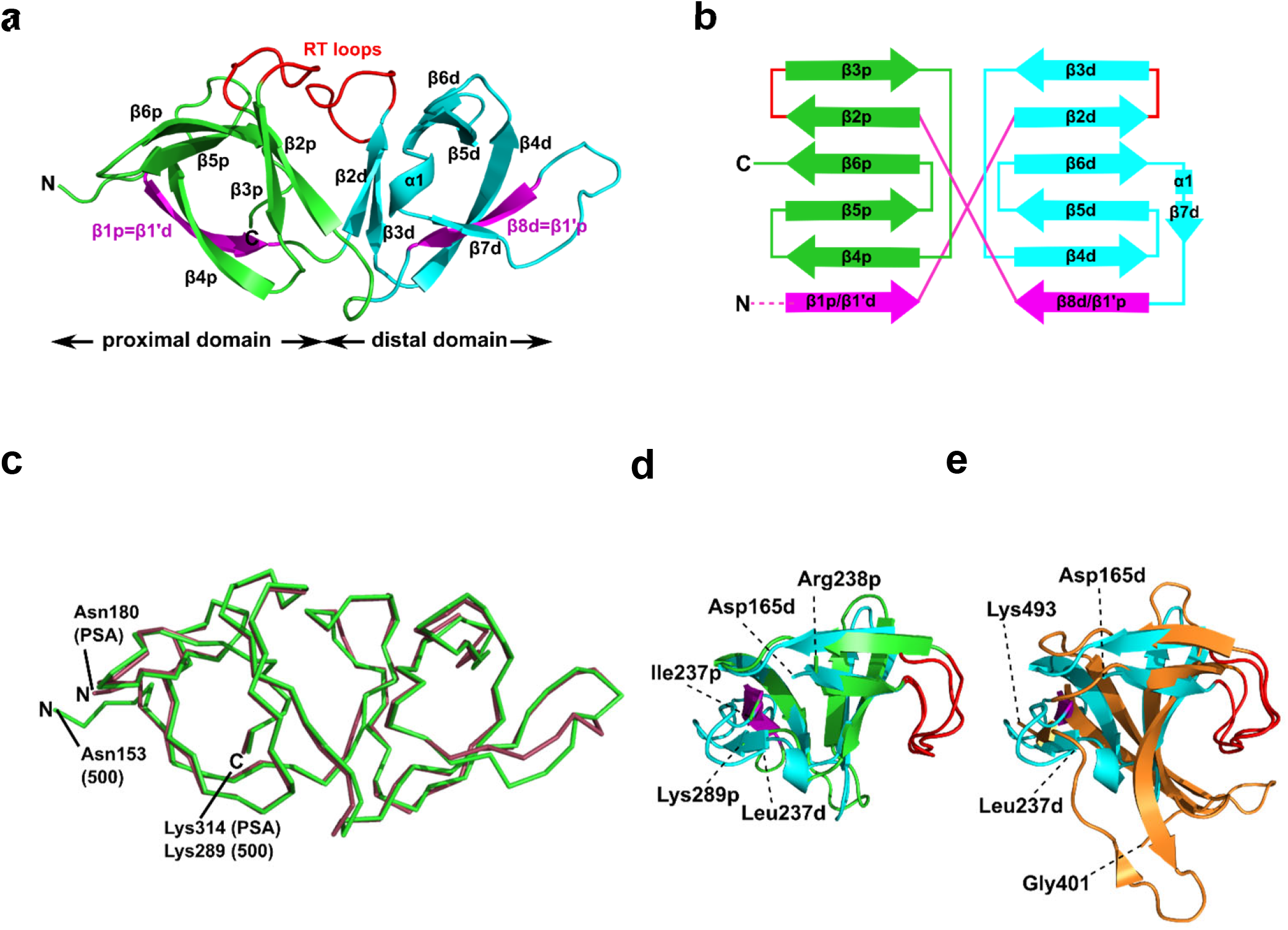
Structural features of CBD500. **a**, Cartoon representation of the proximal (p; green, Cα atoms 153–164 and 238–289) and distal (d; cyan, Cα atoms 165–237) SH3b-like repeats of CBD500 (number corresponds to the full length Ply500). The β-strands β1p and β8d (magenta) are structurally swapped between the two repeats, thus corresponding to β1’d and β1’p, respectively. Loops connecting β2p-β3p and β2d-β3d closely resemble the ligand-binding RT loops^43^ of canonical SH3 domains (red). **b**, Topology diagram of CBD500 with secondary structure elements shown as cylinders for α-helices and arrows for β-strands. **c**, Wire representation of CBD500 and CBDPSA (PDB ID: 1XOV, Cα atoms 180–314) structures upon alignment (RMSD of 0.39 Å for 120 Cα atoms). **d**, Superposition of the pseudo-symmetric (proximal and distal) repeats of CBD500 (RMSD of 1.23 Å, for 34 Cα atoms). **e**, Superposition of the distal repeat of CBD500 (cyan) and the single SH3b domain of lysostaphin calculated by the DALI server^37^ (PDB ID: 5LEO; RMSD of 1.7 Å, Cα atoms 401-493, orange) with the RT loops colored red.

A DALI search^37^ of the PDB with full-length CBD500 identified the CBD of PlyPSA^27^ as a structural homolog (RMSD 0.9 Å for 135 residues), which features the same structurally swapped β-strands connecting its SH3b repeats (**Fig. 3c**). Interestingly, as individual repeats, the proximal and distal subdomains of CBD500 feature high structural similarity (**Fig. 3d**), despite only 17% sequence identity. Both domains also present high structural similarity to s*taphylococcal* bacteriocin CBDs consisting of single SH3b domains, such as ALE-1 (PDB ID: 1R77; RMSD 2.0 Å for 54 Cα atoms)^38^ and Lysostaphin (PDB ID: 5LEO; RMSD 2.1 Å for 58 Cα atoms)^35^, including similar positioning of their individual RT loops (**Fig. 3e**).

### Identification of the WTA-binding site in CBD500 by blind docking and site-directed mutagenesis

To precisely identify the WTA binding site, the 3-O-acetylated repeating unit of WTA_1020_ (GlcNAc-(β1→2)-Rbo-1-PO_4_) was docked onto the crystal structure of CBD500. Three putative cavities with hydrophobic patches were selected. Two of them were also identified by a blind docking carried out with AutoDock Vina^39^ using a grid box comprising the whole CBD500 surface (**Fig. 4a**), with cavity 1 preferentially occupied by ligand. A panel of alanine-scanning mutants within residues of these three cavities were produced and tagged with GFP, to enable quantitative determination of their relative binding capacity to the WSLC 1042 *Listeria* surface by fluorescence spectroscopy (**see Methods**). Changing of W198, W242 and N248 to alanine almost completely abolished the binding, closely followed by W254A, F175A and G250A mutants (**Fig. 4b**). Other mutants showing a significant decrease in bacterial surface binding were Y197A, Y211A and S213A, thus inferring a contribution of the mutated residues to either conservation of the protein conformation or the complex formation. All affected residues are within the cavity 1, confirming the output prediction from the docking study.

**Figure 4.**
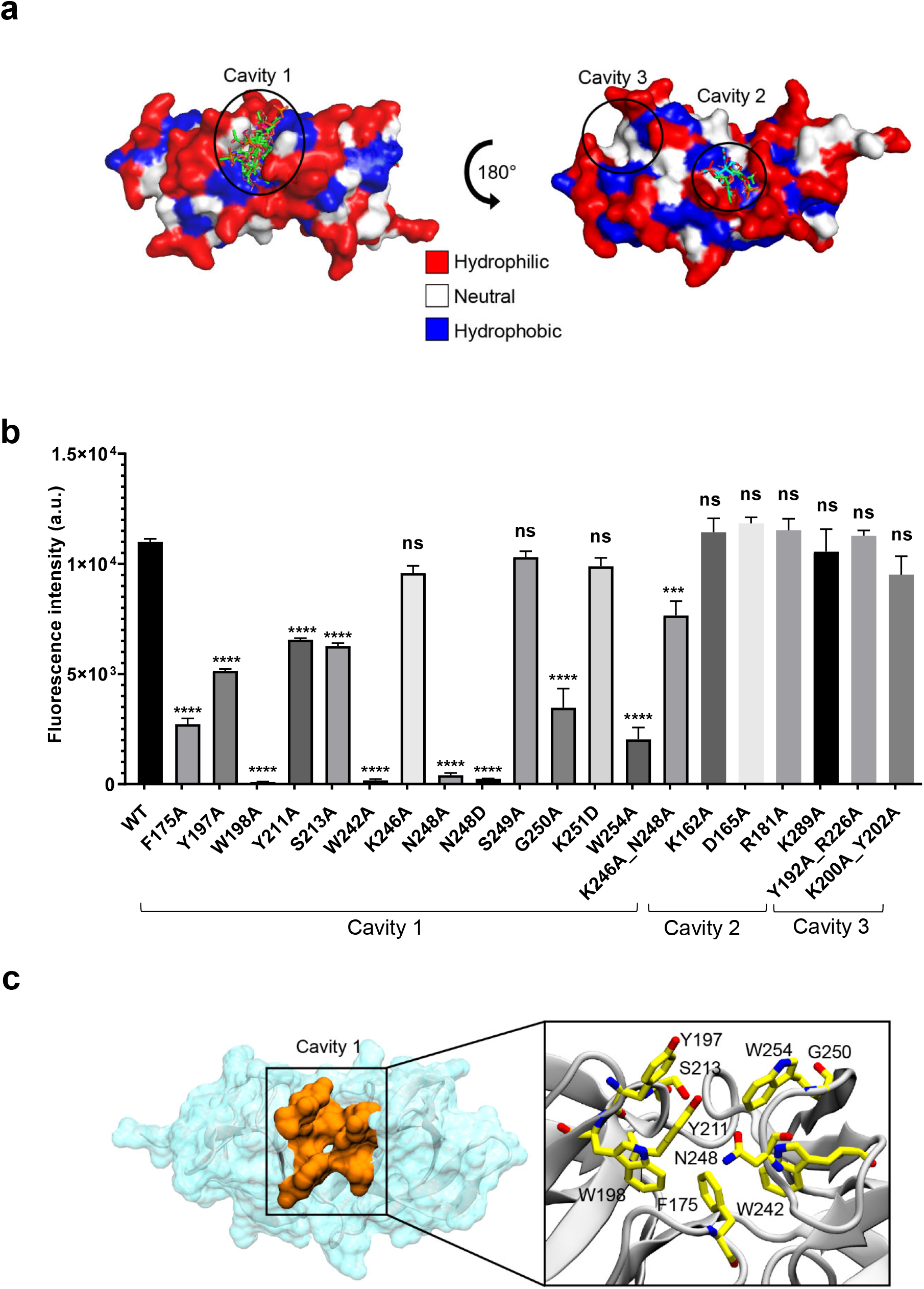
Identification of the ligand binding cavity by blind docking and site-directed mutagenesis. **a**, Putative binding cavities on the surface of CBD500 predicted from blind docking analysis. All ligand poses identified through blind docking are represented in sticks. The CBD500 surface is colored by residue hydrophobicity: hydrophilic (red), neutral (white), and hydrophobic (blue) regions. **b**, Relative binding of GFP-CBD500 mutants to *Listeria* 1042 cells compared to WT, quantified by fluorescence spectroscopy. Results are the mean ± SEM of three experiments with different protein batches. Statistical significance was assessed using one-way ANOVA test (***p<0.001; ****p<0.0001; ns: not significant). **c**, Close-up view of the CBD500 binding cavity 1 (orange). The enlarged view of the square region, shown on the left, depicts the residues (stick representation) shown to be important for binding of CBD500 to *Listeria* surfaces, based on mutational studies.

The molecular weight, chain integrity, and conservation of the protein fold for all GFP-CBD500 mutants were confirmed by SDS-PAGE, analytic gel filtration, and circular dichroism spectroscopy, respectively (**Supplementary Fig. 3a-c**). Only in the case of N248A a size reduction was observed, consistent with a 20 aa truncation at C-terminus identified by MS peptide mapping (data not shown). Another mutant (N248D) did not reveal any sign of truncation, yet still exhibited abolished binding (**Fig. 3b**). Interestingly, the double mutant (K246A_N248A) was able to restore the binding, albeit not to WT level. Furthermore, to reduce the influence of GFP tag, we examined the conformation of several non-tagged CBD500 mutants by analytical gel filtration. The elution profiles of all the mutants were similar to that of the WT (**Supplementary Fig. 3d,e**).

We next tested if the lack of cell wall recognition by select mutants (W198A, W242A, N248A, and W254A) had an effect on the lytic ability of full-length Ply500 endolysin. Ply500 mutants were purified and tested against purified cell wall substrate (**Supplementary Fig. S3f,g**). As expected, all Ply500 mutants bearing an inactivated CBD showed significantly reduced activity^18^, confirming the importance of host recognition by the CBD and of the residues within cavity 1 for modulating the activity of Ply500.

Collectively, the site-directed mutagenesis experiments identified a tweezer-like site in the cavity 1 that possesses a distinctive patch of several hydrophobic and polar residues adjacent to the two RT-loops (**Fig. 4c**).

### Modelling of 3’ *O*Ac-βGlcNAc-Rbo-PO_4_ in complex with CBD500

To identify the feasible interactions between CBD500 and WTA upon complex formation, STD-guided Molecular Dynamics (MD) simulations were run. First, the structure of each complex (CBD500:WTA_1020_, CBD500:WTA_1042-I_ and CBD500:WTA_1042-III_) was modelled by docking of the corresponding 3’ OAc-bearing ligands onto the cavity 1 of CBD500. The MD starting structures were selected based on a combined criteria including favorable binding energy estimation, chain continuity, as well as best fitting between the experimental STD and the theoretically calculated values by applying CORCEMA-ST analysis (**see Methods**)^40^.

For the WTA_1020_ monomer, the CORCEMA-ST analysis pointed towards a single docking conformation for *β*-GlcNAc-(1→2)-Rbo in line with experimental STD values (**Fig. 2a**). This docking output was used as input for MD simulations as described in the **Methods section**. Throughout the simulation the monomeric ligand remained strongly bound at the interface of the RT loops, while the loops and the overall protein structure did not suffer notable distortions (**Fig. 5a**), therefore advocating for the establishment of a single WTA_1020_ repeating unit as the minimal epitope recognized by CBD500. The complete MD 5000-frame trajectory of WTA1020 was subjected to CORCEMA-ST analysis. The profile of the mean theoretical STD values was found to be analogous to the absolute STD experimental values (**Fig. 5b**), further supporting the use of a single repeating unit in the complex modelling. Protons that received the highest saturation transfer were the methyl protons of *O*- and *N*-acetyl groups of 3’ *O*Ac-GlcNAc moiety, followed by protons H3’ and H5’ of the sugar ring (**Fig. 5b**). The MD output was screened for experimentally conformist poses using CORCEMA-ST, yielding two representative monomeric structures for which the calculated absolute STD profile of the sugar ring protons reproduced experimental mean values (**Fig 5b**). The position of the sugar ring remains unchanged at the binding cavity surrounded by several tryptophan residues (**Fig. 5c)**. Both selected structures, belonging to the same and most populated group generated by geographical clustering (see **Methods section**), were found to occupy a position close to the centroid of the cluster (**Fig. 5d**). Even considering with high confidence the GlcNAc ring anchored through the two acetyl groups, the Rbo protons exhibit significant motility throughout the MD (**Fig. 5b-c**), which could be attributed at least in part to the inherent flexibility of Rbo, as expected for a monomeric ligand with open edges.

**Figure 5.**
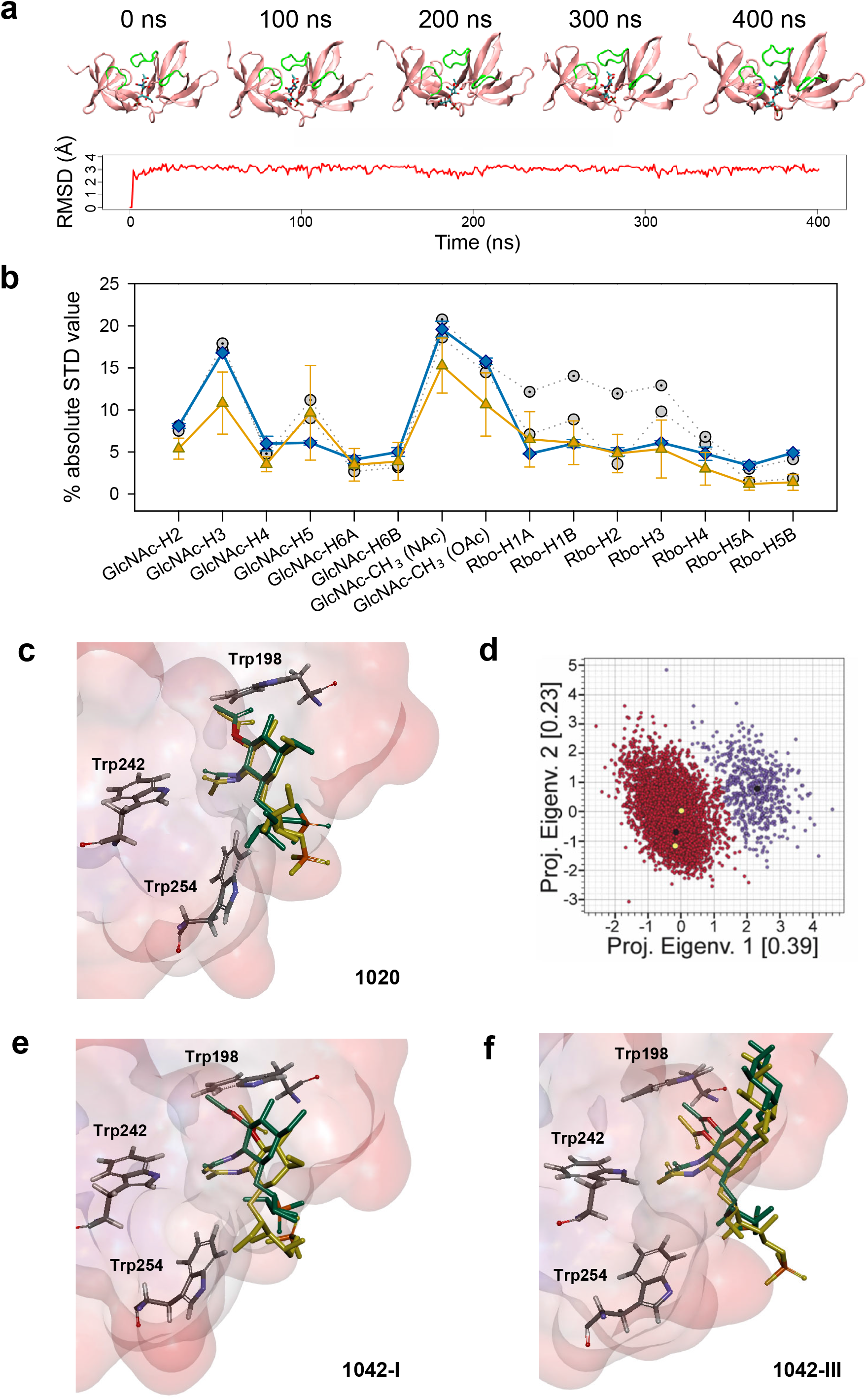
STD NMR-driven MD simulations and structure selection for complex modeling. **a**, Time evolution of the complex of CBD500 with the 3’OAc-bearing WTA_1020_ repeating unit by MD simulations. Representative time-frames (top) and root mean square deviation of the monomeric ligand (bottom). The WTA monomer is represented in sticks, and the CBD500 loops that conform the binding pocket are colored green. **b**, Absolute STD values per proton for the recognized WTA_1020_ repeating unit. The mean ± SD values per proton for the 5000 structures (80ps intervals), extracted from the MD trajectory (orange) and calculated using CORCEMA, is plotted along with the average experimental values from three independent STD-NMR experiments (blue) and the calculated values for the two structures selected on the basis of best correlation to the experimental results (grey). **c**, Superposition of the two selected structures for the CBD500:WTA_1020_ monomer complex. The ligand and relevant Trp residues for reference are represented in sticks and the CBD500 binding pocket in the surface hydrophobicity mode (scale as in **Fig. 4**, panel a). **d**, Eigenvalues Projection map of the 5000 structures (80ps intervals) extracted from the 400 ns MD trajectory of WTA_1020_. The two identified clusters are represented in red and violet respectively, and the black dots represent their centroids. The yellow dots represent the complex structures that were selected on the basis of correlation with experimental results. **e, f**, Superposition of the two selected structures for the CBD500:WTA_1042-I_ and CBD500:WTA_1042-III_ complexes (mode A), respectively. Ligand and protein representation as in **c**.

Docking of WTA_1042_ I and III monomers onto CBD500 yielded two feasible solutions (modes A and B) per complex. Notably, they were up-down opposite orientations (**Supplementary Fig. 4a,b**), where both acetyl groups of 2’-NAc and 3’-OAc of the GlcNAc exchange their relative positions, while still pointing inside the cavity, thus resulting in an inverted orientation of the polymer chain. MD simulations were run for mode A, with the equivalent orientation to the WTA_1020_ monomer pose. Although the ligand remained strongly bound throughout the MD trajectory, the mean theoretical STD profile for both WTA_1042_ monomers did not totally fit with the experimental values obtained with the WT polymer (**Supplementary Fig. 4c,d**). While the calculation predicts the highest saturations for both, *N*- and *O*-acetyl proton groups, larger deviations originate from the calculated values for H4’ and H5’ protons in the opposite faces of GlcNAc pyranose ring, in contrast with MD results of WTA_1020_. As done for WTA_1020_, two representative structures for each monomer of WTA_1042_ I and III were selected after the clustering analysis. After superimposition into the binding site of CBD500, increased motility was observed for both WTA_1042_ monomers compared to that of WTA_1020_ (**Fig 5e,f**). In fact, the two solutions of WTA_1042-I_ showed a considerable shift of the GlcNAc ring, which was significantly reduced by the presence of Gal at position 6’ in WTA_1042-III_. In the context of a polymeric chain, the longer monomer chain and increased rotational freedom provided by ribitol in *β*GlcNAc-(1→4)-Rbo backboned WTAs could also account for a differential response in their recognition by CBD500, compared to *β*(1→2) in WTA_1020_.

### Thermodynamics of the interaction of CBD500 and mutants with WTA polymers

Next, the binding of CBD500 WT and selected mutants to WTA polymers was examined (**Fig. 6a)** by isothermal titration calorimetry (ITC) with emphasis on WTA_1042_ to investigate the binding energetics of complex formation. As often in carbohydrate-protein interactions^41^, the complex formation takes place with a favorable enthalpy change, partially compensated by unfavorable entropic contributions. The affinity of CBD500 for WTA1020 is about three times higher than for WTA_1042_, due to its less unfavorable entropy of binding (**Table 1** **and Supplementary Table 6**). In agreement with fluorescence binding studies, those negative-binding mutants practically abolished the interaction with WTA, whereas the combination of K246A and N248A restored the WTA binding capacity lost by N248A. Mutants Y197A, Y211A, K246A_N248A, and K289A showed a different behavior, as both the enthalpy and the entropy favored their binding to WTA_1042_ and/or WTA1020, indicative of a reduction of hydrogen bonds and/or van der Waals interactions. Remarkably, the stoichiometry of binding for CBD500 WT and the positive-binding mutants was comparable, with approximately 50-60% of all available 3’OAc-GlcNAc units in a WTA polymer chains interacting with a CBD monomer each, and making feasible an avidity increase due to the high local concentration of WTA chains at the bacterial wall surface.

**Figure 6.**
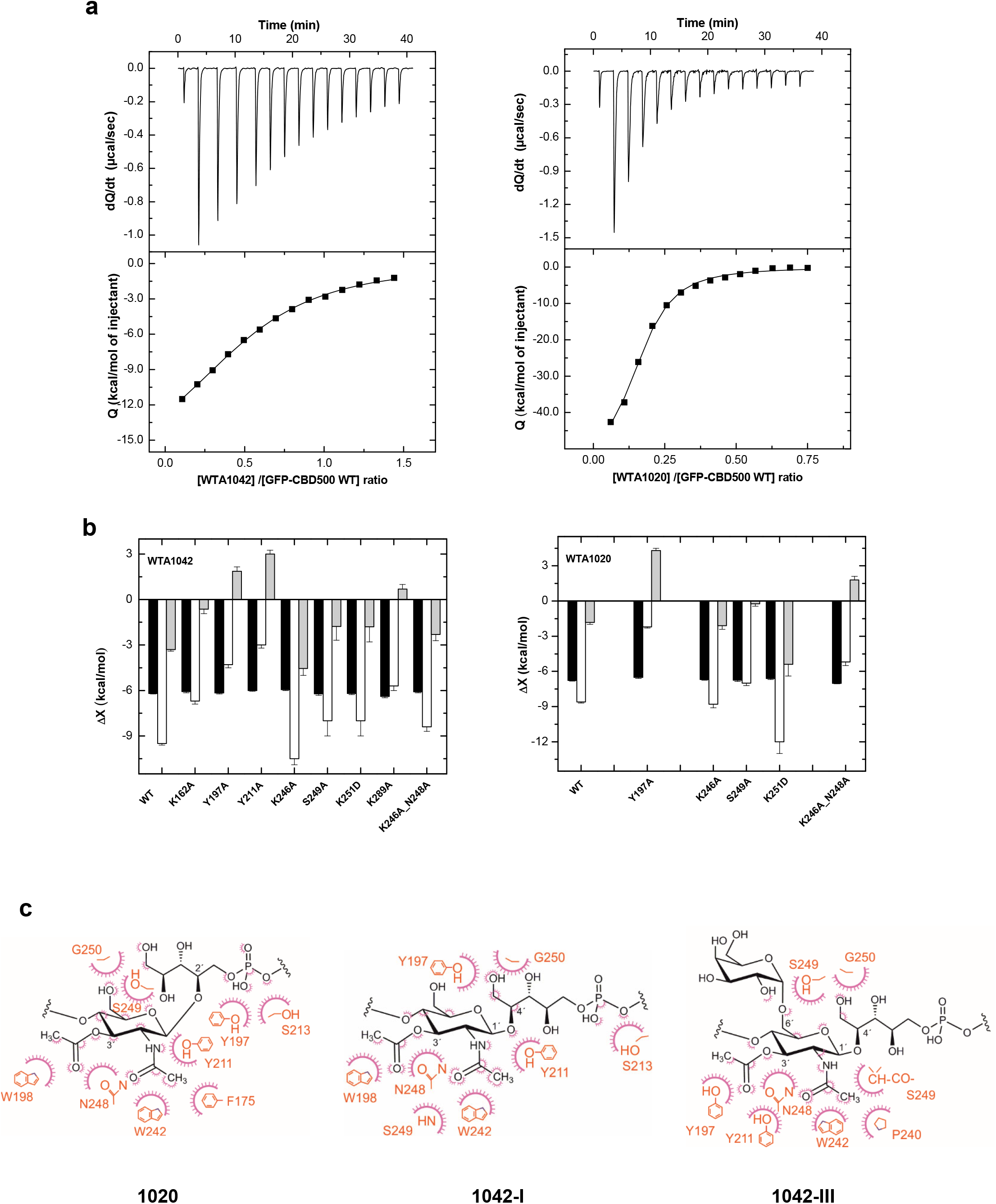
Binding thermodynamics of CBD500 variants to WTA_1020_ and WTA_1042_ and models of complexes. **a**, ITC titration curves of wild-type GFP-CBD500 with WTA_1042_ (left panels) and WTA_1020_ (right panels). Top panels show the raw data, and bottom panels the dependence of the heat evolved per mol of ligand injected with the ligand/protein molar ratio. Solid lines correspond to the theoretical fit of the experimental data (■). See Methods for details. **b**, Thermodynamic dissection of the binding energetics (ΔG = ΔH − TΔS) of wild-type GFP-CBD500 and its active-binding mutants to WTA_1042_ (left) and WTA_1020_ (right). Black bars, ΔG; white bars, ΔH; grey bars, TΔS. **c**, Feasible key non hydrogen-bonded contacts of CBD500 with 3’ *O*Ac-*β*GlcNAc-Rbo-PO4 as derived from the MD analyses (1020 in left panel, 1042-I middle panel and 1042-III right panel). Contacts below 4Å are indicated schematically; the amino-acid residue (in orange) contacts are represented by a purple arc with spokes radiating towards the ligand atoms they contact. The contacted ligand atoms are shown with purple spokes radiating back.

### Atomic interaction of CBD500 with 3’ *O*Ac-*β*GlcNAc-Rbo-PO_4_

The models derived by the STD-driven MD studies for the three feasible complexes (CBD500:WTA_1020_, CBD500:WTA_1042-I_ and CBD500:WTA_1042-III_) displayed both *N*- and *O*-acetyl groups of the GlcNAc unit deeply docked into the binding site, whereas the Gal unit (WTA_1042-III_) sat at a shallower location, near the CBD500 surface. Notably, the amino acid residues inferred, from the functional and thermodynamic data, to be involved in GlcNAc recognition were well conserved in the final selected models based in STD data and MD clustering analysis based on topographical criteria (**Supplementary Fig. 5a-c and Supplementary Table 7**). The GlcNAc ring was consistently involved in bidentate hydrogen bonding through atoms of the acetyl groups, mainly O2N and O1A, and van der Waals contacts mediated by their methyl groups, further supporting the essential role of 3’ -OAc decoration in WTA for CBD500 recognition. Trp242 seemed to establish an undisrupted H-bond with GlcNAc atom O2N through its NE1 proton (RMSD between atoms kept at ~3 Å). Additionally, two main patterns of GlcNAc gripping by Trp198 and Tyr211 were discerned out of the three 400-ns simulations and, remarkably, they appear to be mutually exclusive (**Supplementary Fig. 5b, right panels**). Thus, when Trp198 is hydrogen-bonded to O1A of the 3’ -OAc group, the N2 atom of 2’-NAc group is captured by neighboring Tyr211, as predominantly observed in WTA_1020_ recognition (**Supplementary Fig. 5a**). Positioning of the Rbo-PO_4_ chain in these cases also favored the establishment of H-bonds between the side chain of Ser213 and phosphate oxygen atoms. Alternatively, GlcNAc:O1A can act as acceptor of the OH group of Tyr211 instead of Trp198:NE1, while the distance between Ser213 and the phosphate sharply increases. This pattern was primarily observed throughout the simulations run with WTA_1042_ (**Supplementary Fig. 5b-c and Table 7**). Out of the two WTA_1042-I_ poses selected on the basis of correlation to the experimental STD values for the Δ*gttA* polymer (lacking 6’ Gal, **Fig. 5e**), each showed a distinct recognition pattern – GlcNAc:N2 to Tyr211:OH or Tyr211:OH to GlcNAc:O1A (**Supplementary Fig. 5b**, upper and lower structures respectively) and belonged to separate, comparably populated clusters (data not shown). In contrast, the two selected poses of WTA_1042-III_ (**Fig. 5f**) belonged to the same and most populated cluster (data not shown), and presented the same H-bond pattern with Tyr211:OH acting as donor (**Supplementary Fig. 5c**).

Apart from the aforementioned key residues, additional amino acids participated in an intricate H bond network (**Supplementary Fig. 5b, right panel**), such as Ser249, whose main chain gripped either GlcNAc:O2N (WTA_1042_) or Rbo:HO-5 (WTA_1020_). The neighboring amino acid, Gly250, frequently captured Rbo:O5 in all repeating units, yet atom distance showed high deviation (**Supplementary Table 7**), probably due to the Rbo inherent flexibility. Taken together, the prevalent establishment of a distinct H-bonding pattern in WTA_1042_ compared to WTA_1020_ advocates for a distinct, comprehensive CBD500 recognition mode in a global perspective.

In addition to H bonding, van der Waals interactions^41^ were expected to be particularly relevant in the formation of CBD500-complex (**Fig. 6c**), thanks to methyl-protons and sets of CH-bonds on the ground of a binding site with high density of aromatic residues. MD simulations with the WTA_1020_ monomer also indicated the establishment of van der Waals contacts between the 3’OAc-methyl group of the ligand and the side chain of Asn248, whose loss might also alter the interactions mediated by the Ser249 main chain in the N248A mutant. Phe175, buried in the bottom of the binding groove, was not found to interact directly with WTA_1042_ as with WTA_1020_, but it may play an important structural role through *pi* coordination of aromatic rings. Therefore, our data suggest that both H bonding and van der Waals interactions greatly contribute to the specificity of the carbohydrate-binding CBD500, allowing for ligand selection with structural heterogeneity.

## Discussion

Bacteriophage-encoded endolysins represent a highly attractive class of potent antibacterials, due to their narrow target spectrum, rapid and strong lytic activity. Accordingly, the molecular basis of their cell wall recognition by the endolysin CBD domains is important for development and design of such novel therapeutics. Toward this end, we employed a multidisciplinary strategy, including genetics, biochemical characterization, mutational screening, X-ray crystallography, ligand-based NMR, thermodynamics of binding and computational modelling, to elucidate the structure-function relationship of a bacteriophage endolysin in complex with its WTA ligand.

However, structural studies of WTAs are extremely challenging due to their inherent microheterogeneity and flexibility. Yet, the here described multidisciplinary strategy allows to pinpoint the mechanism by which CBD500 specifically recognizes the 3’ *O*Ac-GlcNAc moiety in the context of a Rbo backbone via a surface-exposed cavity located at the interface of SH3b-like repeats. This cavity comprises multiple aromatic residues that are commonly involved in protein-carbohydrate recognition^42^. The residues located at the cavity make extensive interactions with the two acetyl groups via hydrogen bonds and van der Waals contacts, while permitting the docking of the WTA ligand regardless of the diastereomorphic connectivity (C2 vs. C4) on Rbo and hexose substitution.

We proposed a model for the CBD500-WTA complex (**Supplementary Fig. S6a**), which is new and distinct from current knowledge of ligand interaction with other SH3b-like domains. For instance, the single SH3b of Lysostaphin has been shown to recognize the stem peptide and pentaglycine bridge of PG using a two binding-site mechanism^9^ (**Supplementary Fig. S6b**). The CBD of *Bacillus* endolysin LysPBC15^30^ was shown to bind the peptidoglycan through separate binding sites located on opposite sides of its SH3b domains (**Supplementary Fig. S6c**). Similarly, the CBD of *Clostridium* endolysin PhiSM101^29^ features putative peptide-binding sites facing in opposite directions of ‘pseudo dimer’-like SH3b repeats (**Supplementary Fig, S6d**). This bivalent mode of binding is in strong contrast to the single binding site of CBD500 at the interface between the pseudo-symmetric SH3b repeats. Based on these observations, it appears that the SH3b domains are not homogenous, but have evolved to produce diverse binding sites for recognition of different peptide and carbohydrate ligands. In addition, we found that the SH3b repeats of CBD500 are joined together through β-strand swapping, while the SH3b repeats of both Psm and LysPBC5 fold separately and connect in a more conservative fashion via a short linker. The difference in topology may also contribute to the different mechanisms of ligand binding.

The RT loops appears to be fundamental for binding by SH3 and SH3b domains. In eukaryotic systems, SH3 domains feature a canonical binding site involving the region between the RT and Src loops, which typically binds through protein-protein interactions^43^. In fact, this closely resembles the properties of CBD500 and the lysostaphin SH3b domain that interacts with PG through their RT loops^35^. Despite most *Listeria* phage CBDs feature similar domain architectures, some of them, e.g., CBDP35 and CBD025, appear to recognize different types of WTA polymers^19, 20^. Further research is required to dissect this subtle structural disparity and understand how phage endolysins evolved diverse binding strategies to different ligands at the atomic level. Interestingly, three known *Listeria* virulence factors, InlB, autolysin-like IspC and P60 also feature multiple SH3b repeats at their C-terminus, which have been demonstrated to adhere to glycosyl-polyol teichoic acids^31, 44, 45^. Our data may provide guidance on the target specificity and support future studies of the role of other SH3b-like repeats in virulence and immune defense.

In addition to SH3b, other modules found in the CBDs of endolysins include the LysM domain that binds the PG disaccharide β-*N*-acetylmuramic acid (1 → 4)-β-*N*-acetylglucosamine^46^. The three tandem α-helices repeats of Cpl-7 that recognize the MurNAc residue of PG^8^; the CBDs of Bacillus phage endolysin PlyL and PlyG that binds to the secondary cell wall polysaccharides^47^; and the choline binding repeats described in *Streptococcus pneumonia*e PG hydrolases that interact with the phosphorylcholine residues on teichoic acids through a left-handed *β*-solenoid domain^48^. These variable cell-wall binding modules provide a pool of scaffolds that may be engineered as molecular tools or payloads to target specific bacteria for diagnostic and therapeutic applications.

*O*-acetylation of the MurNAc residues in PG is known to modulate cell wall biosynthesis in most Gram-positive bacteria, and confer resistance against endogenous autolysins and lysozymes of innate immunity systems^49^. However, the presence of *O*-acetylation on other cell wall components has only been uncovered recently, such as in *Listeria* WTA^*20,50*^, *Mesorhizobium loti* LPS^51^, and the GlcNAc residue of *Lactobacillus plantarum* PG^52^. These findings underline the important roles of *O*-acetylation in modulation of autolysin activity, interaction with protozoans, and even virulence^31^. Interestingly, *O*-acetylation of other bacterial surface polysaccharides has also been identified as essential modifications for antibody recognition, and thus needs to be considered in vaccine development^53, 54^. Here, we identified another eminent function of *O*-acetylation of WTA in the recognition by phage endolysins, which are important for phage-bacteria interaction. Surprisingly, the binding affinity (K_D_=*ca*.10^−4^ M measured in solution by ITC analysis) appears significantly lower than the previously reported nanomolar affinities determined by immobilizing whole cells on the SPR surfaces^10^. It is conceivable that weaker local biomolecular interaction may allow the endolysins to better diffuse within the cell wall based on higher dissociation and re-association, and results in an increased avidity through WTA multivalent presentation at cell surface, thereby potentiating the hydrolysis of PG bonds. An intriguing hypothesis is that the mutual adaptability between the binding site and the ligand not only determines the binding specificity, but also boosts the activity of endolysin. The atomic insights and dynamic structures shown in this study will pave new ways towards research in this direction.

## Acknowledgements

We thank Prof. Raffaele Mezzenga and Michael Diener (ETH Zurich) for providing access to circular dichroism spectrophotometers, and are grateful to the staff at the beamline X06SA of the Swiss Light Source (Paul Scherrer Institute, Villigen, Switzerland). E.T.S. has been supported by the Swiss National Science Foundation (SNF) grant 310030_156947/1. FJC and MM have been supported by the Spanish Ministry of Science Innovation and Universities and FEDER funding (grants RTI2018-094751-B-C22 and BFU2015-70052R) and CIBERES, an initiative from the Spanish Institute of Health Carlos III. We acknowledge the technical support by Noelia Hernandez at IQFR and the access to NMR facility at CIB. We thank Sylvain Träger (EPFL) for providing the clustering algorithm and for fruitful discussions.

## Author Contributions

Conceptualization, Y.S., I.K., A.P., M.M., M.D.P., F.J.C., and M.J.L.; Methodology, Y.S., I.K., A.P., M.D. S.B. and M.M.; Investigation, Y.S., I.K., A.P., M.D., S.B., M.M., N.M.I.T., E.T.S., M.R.E., R.M., and A.J.R.; Writing—Original Draft, Y.S., and I.K.; Writing—Review & Editing, Y.S., I.K., A.P. M.D., M.M., M.D.P., F.J.C., and M.J.L.; Visualization, Y.S., I.K., A.P., M.D. S.B., B.G. and M.M; Resources, M.M, P.L. M.D.P., F.J.C., and M.J.L.; Funding Acquisition, M.M., M.D.P., F.J.C., and M.J.L.; Project Administration, Y.S.; Supervision, Y.S., I.K., M.M., M.D.P., F.J.C., and M.J.L.

## Data Availability

The datasets generated and/or analyzed during the current study are available from the corresponding author on reasonable request.

