## Supplemental methods tables figures for "Molecular basis for recognition of *Listeria* cell wall teichoic acid by the pseudo-symmetric SH3b-like repeats of a bacteriophage endolysin"

### Supplementary materials (Shen et al.)

#### METHODS

**Bacterial strains and growth conditions.** All *Listeria* strains were routinely grown in brain heart infusion (BHI) medium with shaking at 30 °C. The deletion mutants were constructed in a *Listeria monocytogenes* serovar 4b strain WSLC1042 background via allelic exchange as previously described[1]. *E. coli* XL1-Blue (Stratagene) used for molecular cloning and protein expression was routinely cultured in Luria–Bertani (LB) broth with shaking at 37 °C.

**Molecular cloning and protein purification.** The plasmid constructs for pQE30::*ply500*, pQE30::*cbd500*, pHGFP::*cbd500* were previously described[2]. The alanine scanning mutant genes of *cbd500* including desired restriction sites for SacI and SalI was commercially synthesized (Thermo-Fischer Scientific). The mutant sequences were subcloned into the pQE30 expression vectors. Purification of His-tagged full length Ply500, C-terminal CBD500, GFP-CBD500 and their variants fused to green fluorescence protein (GFP-tagged) was carried out as previously described[3,4]. The purity of proteins was evaluated by SDS-PAGE. Protein concentration was determined by measuring UV absorbance at 280 nm and using representative extinction coefficients $E_{1\ \mathrm{cm}}^{0.1\%}$ of 2.72 for CBD500, 1.6 for GFP-CBD500, and 2.8 for Ply500, as calculated from their amino acid sequences by the ProtParam tool (http://www.expasy.ch/tools/protparam.html). Selected mutants were further characterized by analytical gel filtration as previously described[5].

**Spot-on-lawn assay.** Overnight cultures of different *Listeria* strains were diluted to an $OD_{600}$ = 0.1 ($l$ = 1 cm) using PBS, and 5 mL of the suspension was spread on 1/2 BHI agar plates. Liquid excess was removed and the plates were air dried for 15 min in a laminar flow hood. One hundred µg $ml^{-1}$ of purified Ply500 endolysins (10 µl) were spotted onto the plates. The plates were air dried for 10 min and then incubated overnight at 37°C. Cleared spots within the resulting bacterial lawn indicate bacterial lysis.

**Dye-release assay.** Cell wall substrates extracted from WSLC 1042 cultures were prepared as described[6] with minor modifications. Briefly, cells (500 ml) were grown to log phase in BHI, heated at 100 °C for 20 min, and then homogenized in a French Press (SLM Aminco) at 100 Mpa. The lysate was centrifuged (20,000*g*, 30 min), weighed and mixed with 200 mM Remazol Brilliant Blue R dye prepared in 250 mM NaOH, at a concentration of 1 g cell wall pellet per 10 ml of solution. Mixtures were incubated 4 °C for 20 h with gentle rocking. Cell wall substrate was centrifuged at 3,000 *g* for 30 min, washed 5 times with 40 ml distilled water each and resuspended in PBS. Assays were performed by mixing 200 µl of substrate suspension ($OD_{595}$ = 1.0, *l* = 1 cm) with 10 µg of wt Ply500 or its variants. Triplicates were prepared for each condition and incubated at 37 °C for 30 min. Cell-wall debris was pelleted by centrifugation at 13,000*g* for 5 min and the $OD_{595}$ of supernatant (100 µl) from each reaction mixture was measured in 96-well flat-bottom plates.

**Lytic killing assay.** *Listeria* cells were grown in BHI medium to mid-log phase, resuspended in PBS at a concentration of $10^7$ CFU $ml^{-1}$. One hundred µl of cells were then mixed with 100 µl of 10 µg $ml^{-1}$ Ply500. PBS alone was used as the negative control. Samples were incubated at 37 °C for 30 min, and the viable counts were performed by serial dilution plating. All assays were performed in triplicate.

**Microscopic visualization of GFP-CBD500 binding to *Listeria* cells.** The fluorescence of GFP-tagged CBD500 (GFP-CBD500) and its variants bound to *Listeria* cell surfaces was examined by confocal laser scanning microscopy as previously reported [7]. Briefly, cells from mid-log phase cultures ($OD_{600}$ = ~0.5) were harvested by centrifugation (10,000*g*, 1 min), and resuspended in PBS (pH 7.4). One hundred µl of cells were incubated with 5 µL of 1 mg $ml^{-1}$ of GFP-CBD500 or its variant at room temperature for 5 min. The cells were washed twice and then resuspended in PBS buffer for confocal laser scanning microscopy. The fluorescence binding assays were performed in triplicate and repeated three times.

**Quantitative evaluation of the binding of GFP-CBD500 and its variants to *Listeria* by fluorescence spectroscopy.** Mid-log phase *Listeria* cells were centrifuged, washed twice, and then resuspended in PBS. Density of bacterial suspension was adjusted to $OD_{600}$ = 1. One hundred µl of cells were incubated with 10 µl of 1 mg $ml^{-1}$ of GFP-CBD500 or its variant, and incubated for 15 min at room temperature. Then, the cells were washed twice and resuspended in PBS buffer (100 µl). Thirty µl of the samples were pipetted into a 96-well black, flat-bottom half area microplate (Greiner Bio-One) and the fluorescence intensity of bound proteins was measured at room temperature using a POLARStar Omega spectrophotometer (BMG Labtech), at 485 nm excitation and 520 nm emission wavelengths with a digital gain of 1,200.

**Extraction and purification of *Listeria* WTA polymers and monomers.** WTA polymers from *Listeria* WSLC 1042 wt and mutant strains were extracted and purified as previously described[7]. The HF-depolymerized products were subject to size exclusion chromatography (Superdex 200 10/300 GL, GE healthcare) in distilled water at 25°C. The corresponding fractions (smaller than 1 kDa) were identified (UV detection at 205 nm), collected and pooled for dialysis (MWCO 100-500 Da, Spectra laboratories) against 1 L of distilled water twice, 12 h each. The dialyzed samples were then lyophilized and stored at - 20 °C for further analysis.

**Ultra-high performance liquid chromatography tandem mass spectrometry (UPLC-MS/MS).** The purified WTA monomers were subjected to UPLC-MS/MS for compositional and structural analysis as previously described[8]. All data were collected and processed using MassLynx software, version 4.1 (Waters), and MS spectra were background-corrected by subtracting the signals between 0–1 min of their respective chromatograms. The relative degree of *O*-acetylation was determined by the ratio:

$$degree\ of\ O\text{-}acetylation\ [\%] = \frac{area(\text{sum of all } O\text{-acetylated peaks})}{area(\text{sum of all peaks})} \qquad \text{(Eq.1)}$$

Similarly, the relative abundance of each isoform in 1042 was dertermined by the ratio

$$relative\ abundancy\ of\ isoform\ X\ [\%] = \frac{area(\text{isoform X peak})}{area(\text{sum of all isoform peaks})} \qquad \text{(Eq. 2)}$$

For example, in the WTA of 1042, the numerator of Eq.1 contains the areas of *O*-acetylated peaks *m*/*z* 396 and 558, and the denominator peaks *m*/*z* 396, 558, 354, 516 (two peaks), and 678 which correspond to all identified repeating units in the polymer. Potential differences in ionization efficiency of the involved species were not considered for this rough estimation. A dilution series of samples confirmed that the relative signal ratios remained constant in the range of relevant signal abundances that were observed for the major species in the various samples analyzed (signal heights of ~3e$^5$ for the major species, or areas of ~60,000 arbitrary units; injection volumes were lowered if the value exceeded).

**Surface plasmon resonance (SPR) analysis.** Sensorgrams of binding of purified *Listeria* WTA wt and mutant polymers to immobilized CBD500 were obtained at 25 °C by SPR (GE Healthcare) as previously described[8] with slight modifications. First, purified CBD500 (0.1 mg ml$^{-1}$ in 10mM sodium acetate buffer, pH 5.0) proteins were immobilized on the carboxymethylated surface of a CMD500L chip (Xantec bioanalytics) at 5 µl min$^{-1}$, using the amine coupling procedure according to the manufacturer's manual. In order to measure and substract non-specific binding, a second flow cell was treated in the same manner, but without immobilization of CBD500 proteins. Approximately seven µM of WTA polymers (analyte) were applied to both cells in running buffer (10 mM Bis-Tris, 100 mM NaCl, 3.4 mM EDTA, 0.005% Tween 20, pH 6) at 10 µL min$^{-1}$ at 25 °C. For each WTA variant, association was followed for 180 s and dissociation for 720 s. The surface was regenerated with a 45 s injection of regeneration buffer (1 M NaCl, 50 mM Tris, pH 9) at a flow rate of 10 µl $min^{-1}$. This cycle was repeated after each measurement.

**Circular Dichroism (CD) spectroscopy.** A Chirascan CD Spectrometer (Bio-Logic) was used to record the CD spectra of CBD domains. Freshly prepared GFP-CBD500 or its mutants were adjusted to a concentration of 0.2 mg $ml^{-1}$ in 10mM phosphate-buffer (pH 7.4). The spectra were recorded at 20 °C using a 0.1-cm path length cuvette. Each scan was obtained by recording ellipticities every 1 nm, with a band width of 1 nm in the range from 190 to 260 nm. The raw spectra were plotted using Origin software (OriginLab).

**NMR Spectroscopy and Saturation Transfer Difference NMR (STD-NMR).** All NMR spectra were acquired at 25 °C on a 600 MHz Bruker Avance spectrometer equipped with a cryoprobe and processed with TopSpin 3.0 software (Bruker). NMR samples were prepared in 99.9% deuterium oxide buffer containing 20 mM phosphate (no NaCl added) at pD 7.2 (uncorrected value). The $^{1}H$ and $^{13}C$ NMR spectra of WTA samples were assigned using a combination of TOCSY (TOtal Correlation SpectroscopY), $^{1}H$-$^{13}C$ HSQC (Heteronuclear Single Quantum Coherence spectroscopy) and ROESY (Rotating-frame Overhauser Enhancement SpectroscopY) of standard pulse sequences. Chemical shifts and coupling constants are reported in the **Tables S1** and **S2** and representative spectra in **Fig. S2a-c**. The $^{1}H$ chemical shifts were referenced to the residual water signal (4.78 ppm at 25 °C) using the equation $\delta$ (ppm) = 5.051 − 0.011 × $T^{e}$ (°C)[9]. TOCSY experiments were performed with 70 ms mixing time, and ROESY experiments were recorded in the phase sensitive mode with water presaturation and the spin-lock module was 300 ms.

Diffusion-ordered spectroscopy (DOSY) yielded an estimation of the WTA polymer mean molecular weight and length and assisted the calculation of the molar concentration of repeating units for the STD and ITC experiments, taking into account the HPLC-MS estimates of relative monomer abundance. DOSY experiments were performed using a stimulated-echo sequence incorporating bipolar gradient pulses and a longitudinal eddy current delay (BPP-LED)[10]. The peak region corresponding to the WTA methyl protons (1.95-2.25 ppm) was selected for integration, and analysis of the raw diffusion data by exponential fitting of the selected peak integral decay to Eq. 3:

$$\ln(I/I_0) = -D\gamma^2 g^2 \delta^2 (\Delta - \delta/3) \qquad \text{(Eq. 3)}$$

where $D$ is the diffusion coefficient, "γ" the gyromagnetic ratio of the nuclei detected, Δ the diffusion delay time, "g" and "δ" the gradient pulse amplitude and duration time, respectively, and $I_0$ is the intensity of a peak when "g" equals to zero. Δ and δ are kept constant during the experiment, while "g" changes. Plotting the normalized resonance intensity for the selected peak region (1.95-2.25 ppm) as a function of the gradient strength and after exponential fitting considering the presence of a sole component, the slope of the resulting line affords the value of *D*. The residual HDO signal yielded a logD value of -8.695. By least-squared fitting of the diffusion coefficient data as a function of the molecular weight for a series of pullulan (linear uncharged polysaccharide) fractions of known characteristics, a calibration curve was obtained that allowed estimating the approximate MW of the WTA (**Fig. S2d-e**), essentially as described[11].

STD experiments were performed at a polymer/protein normality ratio of 25:1 (approx.. 750 μM WTA repeating units and 0.75 mM CBD500) following standard pulse sequences (*stddiff*). Selective saturation of the protein was achieved by using a train of 40 Gaussian-shaped pulses of 50-ms each, separated by a 1 ms delay (approx. saturation time of 2 s). The residual HDO signal was suppressed by excitation sculpting gradient (*esgp*), and CBD background signals, obtained from control experiments performed with protein in the absence of ligand, were subtracted. An off-resonance frequency of δ = 100 ppm and on-resonance frequencies of δ = 0.45/7.1 ppm were applied for selective saturation of CBD500 resonances, targeting spectrum regions where no signals were observed for WTAs. To exclude non-specific saturation of ligand protons upon irradiation, control experiments were performed in the absence of protein (reference STD). For the epitope mapping analysis, spectra with on-resonance frequencies in the aromatic region were selected (7.1 ppm), the highest STD signal was taken as 100% and other STD values were normalized with respect to this signal (**Tables S3 & S4**). Data were derived from at least three independent experiments for wild type WTA polymers, and at least two independent experiments for WTA mutants. For the GlcNAc anomeric proton (H1) no values are reported since quantitation was compromised by solvent suppression.

**Crystallization of His-CBD500.** Crystallization screens of purified His-CBD500 (10 mg $ml^{-1}$; 20 mM Tris, 150 mM NaCl, pH 7.4) were performed in 96-well format plates, using the sitting-drop vapor-diffusion method at 20ºC. Single, needle-like crystals formed after 72 hours with 100 mM bis-Tris, 3 M sodium chloride, pH 5.5, crystallization solution (G12, JBScreen JCSG++, Jena Bioscience GmbH, Jena, DE). No optimization was required to generate larger crystals by hanging-drop vapor-diffusion crystallization, using 1 μl protein (10 mg $ml^{-1}$) mixed with 1 μl crystallization solution, against a 1 ml reservoir of crystallization solution.

**Data Collection and Refinement**. The X06SA (PXI) beamline at the Swiss Light Source, Paul Scherrer Institute, Switzerland, equipped with a PILATUS-6 M (DECTRIS Ltd., Switzerland) pixel detector was used for data collection. A single crystal was fished using a 0.2 mm Round LithoLoop (Molecular Dimensions Ltd., Newmarket, UK). The crystal was cryoprotected by quick soaking in the same crystallization solution containing 30% glycerol, and mounted directly into the liquid nitrogen stream at the beamline. A single, high-resolution data set was collected to 1.59 Å resolution at 100 K using a wavelength of 1.00 Å. Data were automatically processed at the beamline using XDS[12] and SCALA[13], and determined to belong to hexagonal space group P64. Subsequent analysis with phenix.xtriage[14] suggested merohedral twinning, with the twin operator (–h,-k,l), and a twin fraction of 0.449. Data were re-integrated using iMosflm[15] with the correct trigonal space group P32 and scaled again using SCALA. Phenix.sculptor was used to optimize the C-terminal 133 residues of PlyPSA (PDB ID: 1XOV) as a search model for subsequent molecular replacement using PHASER[16]. After an initial refinement using Refmac5[17], successive rounds of refinement were performed using phenix.refine and Coot[18] to generate a final structure with an R factor of 14.4% (Rfree = 18.6%). The stereochemistry of the model was verified with MolProbity[19] and contained 97.0% of the residues within the favored region of the Ramachandran plot and no residues in disallowed regions. All structure figures were created using PyMOL Molecular Graphics System, version 1.4.1, Schrödinger, LLC). The structure is deposited in the Protein Data Bank under Accession Code 6HX0.

**CORCEMA-ST analysis.** Complete relaxation and conformational exchange matrix saturation transfer (CORCEMA-ST)[20,21] analysis enabled the experimentally-driven selection, based on STD data, of ligand poses in the models of CBD500:WTA complexes derived from docking and MD simulations. First, all docking outputs were evaluated for carbohydrate conformational features and the viability of chain continuity from GlcNAc position 4’ and the Rbo-$PO_4$-group in the context of a polymer. Structures fulfilling these criteria were next submitted to CORCEMA-ST analysis, and their calculated theoretical STD NMR values were compared to the experimentally observed values for the polymeric WTA, leading to the selection of the input structures for MD analysis. The same strategy was followed for the 5000 structures extracted from the MD simulation trajectories in order to enable the selection of representative poses for model visualization.

MATLAB scripts of CORCEMA-SW3.8[20-22] were applied in consort with in-house modifications to include experimental saturation values in the input PDB files. The input parameters used in all calculations were 2 s saturation time; a 9 Å-cut around the prospective ligand-pocket; direct irradiation on Tyr, Phe, Trp and His aromatic protons (as an approximation to the selected 7.1 ppm experimental irradiation frequency); and the ITC-derived equilibrium constants of 91000 $M^{-1}$ for WTA 1020 and 35000 $M^{-1}$ for WTA 1042. A $k_{on}$ of $10^8$ L $mol^{-1}$ $s^{-1}$ was used, assuming a diffusion-controlled kinetic model. Correlation times ($\tau_c$) of 1 and 12 ns for the monomeric ligand in the free and bound form, respectively, were estimated following an empirical approximation and considering a 21 kDa monomeric mass for CBD500 ($\tau_c$ = $\rho$*MW/2400 in ns; $\rho$, shape constant)[23]. A concentration of 30 μM for CBD500 was used for calculations, and the concentration for the targeted WTA monomers was adjusted on the basis of their relevant abundance in the native polymer appraised from UPLC-MS/MS.

**Computational modelling with molecular docking and molecular dynamics simulations.**

Docking studies were carried out using the crystal structure of the CBD500 domain as the target, and the following repeating units as the binding ligand: $WTA_{1020}$, $WTA_{1042\text{-}II}$ and $WTA_{1042\text{-}III}$. The structures of repeating units were generated within the Glycam Carbohydrate-builder environment[24] and minimized with the OPLS_2005 force field[25].

Docking was performed with the open source docking software AutoDock Vina[26-29] over the whole protein surface, in order to find all the potential binding sites on CBD500 ("blind" docking). The presence of the solvent was simulated by setting a dielectric constant equal to 78. The ionic concentration was set to 0.15 M for sodium ($Na^+$) and chlorine ($Cl^-$) ions mimicking physiologic conditions. The protonation states of protein residues were assigned with PropKa[30].

The 3D structures used as inputs for Molecular Dynamics (MD) simulations, were constituted by the protein-ligand configurations obtained at the docking stage. Poses with better agreement with STD-NMR experimental data were first selected. The system was solvated with about 20,000 water molecules using a TIP3P water model[31] in order to maintain a physiological water density, and ionized with 0.15 M for $Na^+$ and $Cl^-$. 12 extra $Cl^-$ ions were successively added in order to maintain the whole system electrically neutral, and hence avoid a virtually infinite charge during the MD simulations due to the periodic boundary conditions. CBD500 was parameterized using the Charmm36 force field[32], while WTA repeating units were parameterized with SwissParam[33]. Elastic constraints were applied to the two termini of the WTA repeating unit in order to mimic the presence of the rest of the polymer. The system was first minimized with 15,000 steps of steepest descent algorithm and successively equilibrated with 1 ns of MD at constant volume and temperature (310 K), while the pressure was let free to fluctuate. A velocity rescale algorithm[34] was used to couple the system with a thermal bath, while the Coulomb interactions were treated with the Particle Mesh Ewald method[35]. Finally, a production phase of 400 ns at constant pressure and temperature was run. The pressure of the system was maintained constant with a Parrinello-Rahman barostat[36].

**Clustering, atom distance and H-bonding network analysis.** The atom distances were monitored and plotted through the Visual Molecular Dynamics software[37]. The H-bonding network was also calculated through the same software, by evaluating the atomic distances for each frame. Frames were here extracted at a rate of 1 frame/ns.

A hydrogen bond was considered to be present when the distance between donor and acceptor was below the cut-off value of 3.0 Å; and at the same time, the angle formed by the three atoms was between 150 and 180 degrees. Given their importance for sugar binding[38], presence of CH–*pi* interactions was also scrutinized. The distances between the centroids of H1-H3-H5 GlcNAc protons and the *pi*-electron systems of Phe175, Tyr197, Trp198, Tyr211, Trp242 and Trp254 were monitored. A CH–*pi* distance below 4.0 Å and an angle larger than 120º were considered as prerequisites[39].

The clustering was realized through an in-house developed algorithm. In particular, the positions of all atoms of the ligand throughout the trajectory were extracted and aligned, in order to discard all the translational and rotational movements. The coordinates so obtained were inserted into a matrix, with the atom coordinates in the columns, and the frame numbers in the rows. The eigenvalues of this matrix were calculated, and each ligand structure (corresponding to one single frame) was plotted on a graph with the first and second eigenvalue on the X and Y axis, respectively. According to the distribution of the ligand structures, different clusters were identified around the areas of maximum density. For the clustering analysis, a frame rate of 1 frame/80 ps was used, rendering a collection of 5000 structures, further submitted in CORCEMA-ST analysis for the selection of structures compatible with the STD-NMR data for modelling.

**Isothermal titration calorimetry.** The ITC study of CBD500 titration with full-length WTA was carried out at 25 ºC in a Microcal PEAQ-ITC microcalorimeter (Malvern Instruments). The protein was exhaustively dialyzed against 50 mM $Na_2PO_4H$, 100 mM NaCl, pH 7.4, after elution from the affinity chromatography. WTA solutions were prepared in the same buffer at a monomer concentration of 420 μM for $WTA_{1042}$ and 220 μM for $WTA_{1020}$, based on the DOSY-NMR estimates of respective polymeric chain molecular weights (approximately 15 kDa and 18 kDa) and UPLC-MS estimates of relative monomer abundance. Typically, 1 × 0.4 µl followed by 14 x 2.8 µl injections of WTA were performed automatically into the calorimeter cell loaded with the protein construct, wild type or mutant, at a monomer concentration of 10-87 µM, under 750 rpm stirring. Prior to data analysis, the heat contributed by WTA or the protein dilution was measured in separate runs and subtracted, when required, from the total heat produced following each injection during titration. Titration curves were analyzed with Microcal-ITC Origin (ligand-in-cell option) and Affinimeter-ITC softwares, assuming a model of *N* equivalent and independent sites per WTA polymeric chain.

**Statistics.** All experiments were performed with n equal to or greater than 3 unless otherwise stated. Statistical analysis using one-way ANOVA test is reported as *$p<0.05$; **$p<0.01$; ***$p<0.001$; ****$p<0.0001$; ns: not significant.

Supplementary Table 1. $^{1}$H- and $^{13}$C-NMR chemical shifts for WTA 1042 variants (in ppm).

| **Variant** | **Residue** | **Sugar unit** | $^{3}J_{H,H}$† | **H1 (C1)** | **H2 (C2)** | **H3 (C3)** | **H4 (C4)** | **H5 (C5)** | **H6$_A$ (C6)** | **H6$_B$** | **NAc-CH$_3$** | **OAc-CH$_3$** |
|---|---|---|---|---|---|---|---|---|---|---|---|---|
| I-VI[a] | 3´Glc-GlcNAc | [α-Gal]<br>↓ 6´<br>→4)-β-D-GlcNAc-(1→<br>↑3´ Glc | 7.5 | 4.92<br>(100.2) | *4.00*<br>(55.1) | 4.07<br>(78.0) | 4.12<br>(73.2) | 3.60 [3.87]<br>(75.1) [73.8] | *3.82* [*3.89*]<br>(60.4) [66.2] | *3.90* [*3.95*] | 2.08<br>(22.6) | N/A |
| II-V[b] | GlcNAc | [α-Gal]<br>↓ 6´<br>→4)-β-D-GlcNAc-(1→ | 8.2 | 4.65-4.69<br>(101.0) | *3.82*<br>(55.3) | *3.73*<br>(71.8) | *4.00*<br>(74.2) | 3.56 [*3.88*]<br>(75.5)[81.9] | *3.82* [*3.89*]<br>(60.4) [66.2] | *3.90* [*3.95*] | 2.03<br>(22.1) | N/A |
| III-IV | 3´OAc-GlcNAc | [α-Gal]<br>↓ 6´<br>→4)-β-D-GlcNAc-(1→<br>↑3´ OAc | 8.0 | 4.82<br>(100.4) | 3.93 | 5.19<br>(74.3) | 4.21 [4.33]<br>(71.6) [71.8] | 3.67 [*3.90*]<br>(74.7) [73.4] | *3.82* [*3.89*]<br>(60.4) [66.2] | *3.90* [*3.95*] | 2.00-2.02<br>(22.2) | 2.13<br>(20.6) |
| I-VI | Glc | β-D-Glc-(1→ | 7.7 | 4.54<br>(101.9) | *3.34*<br>(72.9) | 3.45<br>(75.5) | 3.44<br>(69.4) | *3.34*<br>(75.5) | 3.71<br>(60.4) | 3.84 | N/A | N/A |
| I-III[c] | Gal | α-D-Gal-(1→ | 3.6 | 5.05<br>(98.3) | 3.81<br>(68.4) | *3.91*<br>(69.4) | 3.98<br>(69.3) | 4.03<br>(71.0) | 3.74<br>(61.2) | | N/A | N/A |
| | | | | **H1$_A$ (C1)** | **H1$_B$** | **H2 (C2)** | **H3 (C3)** | **H4 (C4)** | **H5$_A$ (C5)** | **H5$_B$** | | |
| all | Rbo | →4)-Rbo<br>↑1´ PO4-(→ | | 3.92<br>(66.5) | 4.11 | 3.78<br>(70.2) | 3.83<br>(70.6) | *3.91*<br>(82.0) | *3.73*<br>(60.7) | *3.88* | | |

Spectra were acquired in deuterated phosphate buffer at 298 K and the proton chemical shifts were referenced to the residual HDO signal (4.78 ppm at 25° C). Values given in brackets correspond to the Gal-bearing variants. † The $^{3}J_{H,H}$ values (in Hz) for the anomeric proton resonances (H1) were calculated from TOCSY spectra. [a,b]Assignment of chemical shifts for the variant I was aided by acquired spectra of the *ΔoatT* mutant and for variant II from spectra of the *ΔgalU* at the same experimental conditions and buffer. [c]Applies for galactosylated variants. N/A: not applicable. Peaks overlapping in the $^{1}$H dimension, thus affecting STD quantitation, are given in *italic*.

Supplementary Table 2. $^{1}$H and $^{13}$C chemical shifts for WTA 1020 variants (in ppm).

| **Variant** | **Residue** | **Sugar unit** | $^{3}J_{H,H}$† | **H1<br>(C1)** | **H2<br>(C2)** | **H3<br>(C3)** | **H4<br>(C4)** | **H5<br>(C5)** | **H6$_A$<br>(C6)** | **H6$_B$** | **NAc-$CH_3$** | **OAc-$CH_3$** |
|---|---|---|---|---|---|---|---|---|---|---|---|---|
| I | GlcNAc | →4)-β-D-GlcNAc-(1→ | 8.5 | 4.73<br>(101.4) | 3.79<br>(55.6) | 3.73<br>(73.0) | *3.96*<br>(74.0) | 3.55<br>(74.9) | *3.81*<br>(60.4) | *3.90* | 2.09<br>(22.3) | – |
| II | 3´OAc-GlcNAc | →4)-β-D-GlcNAc-(1→<br>↑3´OAc | 8.2 | 4.87<br>(100.7) | 3.94<br>(54.0) | 5.16<br>(74.9) | *4.18*<br>(71.7) | 3.65<br>(74.8) | *3.81*<br>(60.4) | *3.90* | 2.03<br>(22.1) | 2.13<br>20.6 |
| | | | | **H1$_A$<br>(C1)** | **H1$_B$** | **H2<br>(C2)** | **H3<br>(C3)** | **H4<br>(C4)** | **H5$_A$<br>(C5)** | **H5$_B$** | | |
| all | Rbo | →2)-Rbo<br>↑1´PO4-<br>(→ | | *3.96*<br>(65.3) | *4.18* | 4.10<br>(79.6) | 3.80<br>(71.3) | 3.83<br>(71.8) | 3.62<br>(62.5) | 3.82 | | |

Spectra were acquired in deuterated phosphate buffer at 298 K and the proton chemical shifts were referenced to the residual HDO signal (4.78 ppm). † The $^{3}J_{H,H}$ values (in Hz) for the anomeric proton resonances (H1) of the GlcNAc moieties were calculated from TOCSY spectra. Peaks overlapping in the $^{1}$H dimension, thus affecting STD quantitation, are given in *italic*.

Supplementary Table 3. STD intensities for the CBD500-WTA$_{1020}$ and CBD500-WTA$_{1042}$ complexes.

| WTA | Proton | STD value | Normalized STD value[a] | Proton | STD value | Normalized STD value[a] | Proton | STD value | Normalized STD value[a] |
|---|---|---|---|---|---|---|---|---|---|
| **1020** | **→4)-GlcNAc-(1→ ↑3´OAc** | | | **→2)-Rbo ↑1´ PO$_4$-(→** | | | | | |
| | H1 | N/D | N/D | H1$_A$ | *4.8%* | *24%* | | | |
| | H2 | 8.1% | 41% | H1$_B$ | *6%* | *25%* | | | |
| | H3 | 16.8% | 86% | H2 | 5% | 26% | | | |
| | H4 | *6%* | *31%* | H3 | 6.1% | 31% | | | |
| | H5 | 6.1% | 31% | H4 | 4.8% | 24% | | | |
| | H6$_{A-B}$ | *4.1%* ; *5.1%* | *21%*; *26%* | H5$_A$ | 3.4% | 17% | | | |
| | NAc-CH$_3$ | 19.6% | 100% | H5$_B$ | 4.9% | 25% | | | |
| | OAc-CH$_3$ | 15.8% | 81% | | | | | | |
| **1042** | **[αGal] ↓ 6´ →4)-GlcNAc-(1→ ↑3´ OAc** | | | **→4)-Rbo ↑1´ PO$_4$-(→** | | | **αGal-(1→** | | |
| | H1 | N/D | N/D | H1$_A$ | 4.6% | 18% | H1 | 2.9% | 12% |
| | H2 | 7.5% | 30% | H1$_B$ | 4% | 16% | H2 | 5.8% | 23% |
| | H3 | 16.3% | 64% | H2 | 4.5% | 18% | H3 | *4.1%* | *16%* |
| | H4 | 14.6 [14.8]% | 56 [58]% | H3 | 5.3% | 21% | H4 | 3.3% | 13% |
| | H5 | 4.5 [*3.1*]% | 18 [*12*]% | H4 | *4.1%* | *16%* | H5 | 2.7% | 11% |
| | H6$_{A-B}$ | *4.5* [*3.1*]%; | *18* [*12*]%; | H5$_A$ | *2%* | *8%* | H6$_{A-B}$ | *2.5%* | *10%* |
| | NAc-CH$_3$ | 23.8% | 94% | H5$_B$ | *3.3%* | *13%* | | | |
| | OAc-CH$_3$ | 25.3% | 100% | | | | | | |

The reported absolute STD values represent the mean of at least three independent experiments, registered upon aromatic irradiation (7.1 ppm) of 1:25 CBD:WTA mixtures (30 μM CBD500 and approx. 750 μM WTA monomeric units). Values given in brackets correspond to the Gal-bearing variants. [a]Normalized values were calculated considering the highest transfer signal as 100%. Signals overlapping in the $^1$H dimension, as deduced from the chemical shifts assignment (Suppl. Table 1 and 2), are given in *italic*. The mean STD value is given for these overlapping peaks. N/D: not defined, quantitation is compromised by solvent suppression.

Supplementary Table 4. STD intensities for the CBD500-WTA$_{1042}$Δ*gttA*, Δ*galU* and Δ*gltB* complexes.

| WTA | Proton | STD value | Normalized STD value[a] | Proton | STD value | Normalized STD value[a] |
|---|---|---|---|---|---|---|
| | →4)-GlcNAc-(1→ ↑3´ OAc | | | →4)-Rbo ↑1´ $PO_4$-(→ | | |
| **1042Δ*gttA*** | H1 | N/D | N/D | H1$_A$ | 3.1% | 12% |
| | H2 | 3.1% | 12% | H1$_B$ | 2.8% | 11% |
| | H3 | 14.5% | 55% | H2 | 2.8% | 11% |
| | H4 | 13.1% | 49% | H3 | 3.8% | 14% |
| | H5 | 6.4% | 24% | H4 | 2.7% | 10% |
| | H6$_{A-B}$ | 5.5%; 2.1% | 21%; 8% | H5$_A$ | 1.7% | 6% |
| | NAc-$CH_3$ | 19.2% | 73% | H5$_B$ | 1.9% | 7% |
| | OAc-$CH_3$ | 26.5% | 100% | | | |
| **1042Δ*galU*** | H1 | N/D | N/D | H1$_A$ | 3.6% | 27% |
| | H2 | 4.2% | 31% | H1$_B$ | 2.5% | 19% |
| | H3 | 11.5% | 86% | H2 | 2.9% | 22% |
| | H4 | 8.4% | 63% | H3 | 2.8% | 21% |
| | H5 | 3.7% | 28% | H4 | 3.5% | 26% |
| | H6$_{A-B}$ | 3.6%; 2.3% | 27%; 17% | H5$_A$ | 3.8% | 28% |
| | NAc-$CH_3$ | 10.8% | 81% | H5$_B$ | 2.4% | 18% |
| | OAc-$CH_3$ | 13.4% | 100% | | | |
| | [*α*Gal] ↓ 6´ →4)-GlcNAc-(1→ ↑3´ OAc | | | →4)-Rbo ↑1´ $PO_4$-(→ | | |
| **1042Δ*gltB*** | H1 | N/D | N/D | H1$_A$ | 4.9% | 31% |
| | H2 | 5.1% | 33% | H1$_B$ | 4.5% | 29% |
| | H3 | 14.8% | 95% | H2 | 5.2% | 33% |
| | H4 | 13.1 [12.8]% | 84% [82%] | H3 | 5.3% | 34% |
| | H5 | 3.9 [4.1]% | 25% [26%] | H4 | 4.8% | 31% |
| | H6$^{b}_{A-B}$ | 5.1%; 4.2% | 33%; 27% | H5$_A$ | 2.4% | 15% |
| | NAc-$CH_3$ | 14.3% | 92% | H5$_B$ | 4.1% | 26% |
| | OAc-$CH_3$ | 15.6% | 100% | | | |

The reported absolute STD values represent the mean of at least two independent experiments, registered upon aromatic irradiation (7.1 ppm) of 1:25 CBD:WTA mixtures (30 μM CBD500 and approx. 0.75/0.9/1 mM WTA monomeric units for Δ*gttA*, Δ*gltB* and Δ*galU*, respectively. [a]Normalized values were calculated considering the highest transfer signal as 100%. N/D: not defined, quantitation is compromised by solvent suppression. [b]For this 6'Gal-bearing monomer, no significant associated difference is observed for the STD-NMR values.

Supplementary Table 5. Crystal structure data collection, phasing and refinement statistics.

| **Data Collection** | His-CBD500 (aa 133-289) |
|---|---|
| Space Group | $P3_2$ |
| Cell Dimensions | |
| *a, b, c* (Å) | 49.58, 49.58, 63.77 |
| α, β, γ (°) | 90, 90, 120 |
| Wavelength (Å) | 1.00 |
| Resolution Range (Å) | 63.77 - 1.59 (1.67-1.59)* |
| Number of Unique Reflections | 23,622 |
| $R_{meas}$ | 0.168 (1.073) |
| $I/\sigma I$ | 8.5 (2.4) |
| Completeness (%) | 99.6 (99.4) |
| Redundancy | 9.0 (8.3) |
| Twin Law | -h,-k,l |
| **Refinement** | |
| Resolution (Å) | 42.94 - 1.59 |
| No. Reflections | 23,591 |
| $R_{work}$ / $R_{free}$ | 0.144 / 0.186 |
| No. Atoms | |
| Protein | 1,182 |
| Ligand/Ion | 0 |
| Water | 461 |
| B-factors (Å$^2$) | |
| Protein | 18.7 |
| Ligand/Ion | 0 |
| Water | 36.95 |
| Ramachandran Plot (%) | |
| Favored | 97 |
| Outliers | 0 |
| R.M.S Deviations | |
| Bond Lengths (Å) | 0.006 |
| Bond Angles (°) | 0.67 |

*Highest resolution shell is shown in parenthesis

Supplementary Table 6. *Listeria* binding capacity of GFP-CBD500 alanine mutants and binding thermodynamics to WTA1020 as derived from ITC.

| Constructs | Binding to *Listeria*[a] | N[b] | $K_b$ ($M^{-1}$) | -ΔH (kcal/mol) | -ΔG (kcal/mol) | TΔS (kcal/mol) |
|---|---|---|---|---|---|---|
| WT | + | 6.5±0.1 | (9.1±0.8) ×$10^4$ | 8.6±0.1 | 6.77±0.05 | -1.83±0.15 |
| Y197A | (+) | 4.0±0.3 | (6±1) ×$10^4$ | 2.2±0.2 | 6.5±0.1 | 4.3±0.3 |
| K246A | + | 5.2±0.1 | (8.2±0.4) ×$10^4$ | 8.8±0.3 | 6.70±0.03 | -2.1±0.3 |
| S249A | (+) | 4.5±0.1 | (9±2) ×$10^4$ | 7.0±0.2 | 6.8±0.1 | -0.2±0.3 |
| K251D | + | 4.4±0.5 | (7±1) ×$10^4$ | 12±1 | 6.61±0.08 | -5±1 |
| K246A_N248A | + | 4.7±0.3 | (1.4 ±0.1) ×$10^5$ | 5.2±0.3 | 7.02±0.04 | 1.8±0.3 |
| I252A | (+) | Protein aggregation avoided quantification of ITC data | | | | |
| W254A | (+) | Extremely weak affinity not quantifiable by ITC | | | | |
| N248A[c] | - | Extremely weak affinity not quantifiable by ITC | | | | |
| GFP | - | No binding detected by ITC | | | | |

[a]Binding was visualized by fluorescence microscopy using the GFP-CBD500 construct and its variants: '+', strong binding; '(+)', weak binding; '-', no binding.

[b]N is the number of CBD molecules bound per WTA chain. $K_b$, is the binding constant per monomer of CBD bound to WTA as measured by ITC, and ΔG ($-RT\ln K_b$), ΔH and ΔS represent the corresponding thermodynamic parameters.

[c]ITC measured with both CBD500 and GFP-CBD500 mutants

Supplementary Table 7. Summary of H-bonding network derived from MD simulations of CBD500 in complex with the repeating unit of $WTA_{1020}$, $WTA_{1042\text{-}I}$, and $WTA_{1042\text{-}III}$.

| Complex | Sugar Unit | Sugar Atom | Protein residue : Atom | # of frames where H-bond present | Mean Distance (Å) | # of frames where H-bond not present | Mean Distance (Å) |
|---|---|---|---|---|---|---|---|
| CBD500/$WTA_{1020}$ | GlcNAc | O1A | W198 : NE1 | 72/401 | 2.84 ± 0.10 | 329/401 | 3.67 ± 0.77 |
| | GlcNAc | N2 | Y211 : OH | 169/401 | 2.87 ± 0.07 | 232/401 | 3.29 ± 0.35 |
| | GlcNAc | O2N | W242 : NE1 | 189/401 | 2.85 ± 0.09 | 212/401 | 3.15 ± 0.16 |
| | GlcNAc | O6 | S249 : OG | 23/401 | 2.86 ± 0.08 | 378/401 | 5.14 ± 1.10 |
| | Rbo | O1 | S213 : OG | 109/401 | 2.73 ± 0.10 | 292/401 | 5.09 ± 1.44 |
| | Rbo | O5 | G250 : O | 96/401 | 2.79 ± 0.11 | 305/401 | 5.35 ± 1.63 |
| CBD500/$WTA_{1042\text{-}I}$ | GlcNAc | O1A | W198 : NE1 | 25/401 | 2.86 ± 0.09 | 376/401 | 4.85 ± 1.03 |
| | GlcNAc | N2 | Y211 : OH | 46/401 | 2.87 ± 0.08 | 355/401 | 3.63 ± 0.43 |
| | GlcNAc | O2N | W242 : NE1 | 174/401 | 2.85 ± 0.10 | 227/401 | 3.14 ± 0.12 |
| | GlcNAc | O2N | S249 : N | 157/401 | 2.85 ± 0.09 | 244/401 | 3.21 ± 0.26 |
| | GlcNAc | O1A | Y211 : OH | 143/401 | 2.73 ± 0.11 | 258/401 | 5.01 ± 1.17 |
| | GlcNAc | O5 | S249 : OG | 42/401 | 2.88 ± 0.09 | 359/401 | 3.94 ± 0.95 |
| | GlcNAc | O6 | S249 : OG | 0/401 | NA | 401/401 | 7.00 ± 1.52 |
| | GlcNAc | O3 | N248 : ND2 | 0/401 | NA | 401/401 | 5.86 ± 0.69 |
| | Rbo | O5 | G250 : O | 86/401 | 2.80 ± 0.12 | 315/401 | 4.69 ± 1.13 |
| CBD500/$WTA_{1042\text{-}III}$ | GlcNAc | O1A | W198 : NE1 | 59/401 | 2.86 ± 0.09 | 342/401 | 4.01 ± 0.81 |
| | GlcNAc | N2 | Y211 : OH | 100/401 | 2.88 ± 0.08 | 301/401 | 3.46 ± 0.44 |
| | GlcNAc | O2N | W242 : NE1 | 205/401 | 2.84 ± 0.09 | 196/401 | 3.13 ± 0.14 |
| | GlcNAc | O2N | S249 : N | 107/401 | 2.87 ± 0.09 | 294/401 | 3.22 ± 0.20 |
| | GlcNAc | O1A | Y211 : OH | 111/401 | 2.70 ± 0.10 | 290/401 | 4.72 ± 0.51 |
| | GlcNAc | O5 | S249 : OG | 59/401 | 2.88 ± 0.08 | 342/401 | 3.59 ± 0.49 |
| | Rbo | O1 | S213 : OG | 63/401 | 2.76 ± 0.10 | 338/401 | 5.93 ± 1.73 |
| | Rbo | O5 | G250 : O | 53/401 | 2.82 ± 0.11 | 348/401 | 4.29 ± 0.81 |

List of the hydrogen bonds observed across the 400-ns MD trajectories. The atoms and chemical groups involved, the frequency of observation and the mean distance between the aminoacids are also reported.

### Figure legends

**Figure S1. *Listeria* WTA decoration and its effect on CBD500 binding. a**, Liquid chromatographic separation and MS-based identification of carbohydrate residues within WTAs *Lm* strain WLSC 1042 and its 1042Δ*oatT* mutant, lacking *O*-acetylation. Peaks corresponding to the 3' *O*-acetylated monomers of $WTA_{1042}$ are labeled with base peak ion $[M\text{-}H]^-$ (m/z) values. **b**, Representative, raw SPR sensorgrams of 7 μM WTA polymer (analyte) recognition by immobilized CBD500 out of three individual experiments (RU: response units). WTAs were purified from the wild-type 1042 bacteria or its mutants as indicated. **c**, Degree of *O*-acetylation in the indicated strains, as determined by UPLC-MS/MS analysis using Eq. 1 (Data represent mean ± SEM; n=3). Statistical significance was assessed using one-way ANOVA test (****p<0.0001; ns: not significant). **d**, Binding of GFP-CBD500 to the *Listeria ivanovii* strain WSLC 3009 and its *O*-acetylation deficient mutant, 3009Δ*oatT* (scale bar: 1 μm). **e**, Comparison of UPLC-MS/MS chromatograms of WTA monomers isolated from WSLC 3009 wt and WSLC 3009Δ*oatT* mutant. The peak corresponding to the 3'*O*-acetylated monomers is labeled with its base peak ion $[M\text{-}H]^-$ (m/z).

**Figure S2. NMR characterization of WTA polymers and molecular weight estimation by DOSY NMR. a-c**, Sections of the multiplicity-edited $^1H$-$^{13}C$ HSQC NMR spectra of $WTA_{1020}$ (**a**) and $WTA_{1042}$ (**b**), and of WTA derived from $WTA_{1042}$*ΔgalU* and $WTA_{1042}$*ΔoatT* mutants (**c**). Detailed assignment information is included in **Supplementary Tables 1** and **2**. Signals labelled with an asterisk were not assigned. $CH_2$ protons are shown in cyan, while CH and $CH_3$ are in black ($WTA_{1020}$, $WTA_{1042}$ and $WTA_{1042}$*ΔgalU*) or purple ($WTA_{1042}$*ΔoatT*). In **a**, the inset represents a zoom of the zone of methyl protons at $\delta$ ~2 ppm ($^1H$, F2 axis), while a diffusion-filtered $^1H$ spectrum is overlapped with the $^1H$-$^{13}C$ HSQC, in both **a** and **b**. **d**, The sum of columns of the DOSY spectra (NMR signal versus logD representation) of $WTA_{1042}$ (black) and $WTA_{1020}$ (red) resulted in log*D* values of -10.14 and -10.18, respectively. **e,** Logarithmic plot of MW against log*D* values for eight pullulan fractions of known molecular weight (•). All data points are located on a straight line used for molecular weight calibration. Experimental data points for $WTA_{1042}$ (**x**) and $WTA_{1020}$ (**x**) are shown.

**Figure S3. Biochemical and biophysical characterization of GFP-CBD500, CBD500 and Ply500 variants. a,** SDS-PAGE of selected GFP-CBD500 variants purified from nickel affinity chromatography. **b**, Elution profiles of selected GFP-CBD500 variants in analytical gel filtration chromatography. **c**, Far-UV CD spectra of GFP-CBD500 WT and its mutants. Data are of three individual experiments. **d**, SDS-PAGE of selected CBD500 variants purified by nickel affinity chromatography. **e**, Comparison of CBD500 variant elution profiles in analytical gel filtration chromatography. **f**, SDS-PAGE of selected Ply500 variants purified by nickel affinity chromatography. **g**, Cell wall hydrolytic activity of selected Ply500 mutants on *Listeria* 1042 cells determined by the dye-release colorimetric assay. Results are the mean ± SEM of three separate experiments. Statistical significance is assessed using one way ANOVA test (****p<0.001).

**Figure S4. STD NMR-driven MD simulations and ligand pose selection for modeling the interaction of CBD500 with $WTA_{1042\text{-}I}$ and $WTA_{1042\text{-}III}$ monomers.** Panels **a)** and **b)**: Close-up view of the two initially selected docking output poses in the binding site with relevant Trp residues in stick representation for the $WTA_{1042\text{-}I}$ monomer **a)** and for the $WTA_{1042\text{-}III}$ monomer **b);** Panels **c)** and **d)**: Experimental values for the polymeric $WTA_{1042}$ wild type (n=3, blue) plotted together with CORCEMA-calculated absolute STD values per GlcNAc proton for the WTA monomers from the MD-extracted 5000 structures (mean ± SD, orange) for **c)** the $WTA_{1042\text{-}I}$ monomer and the two selected structures shown in **Fig. 5e** (grey), and **d)** the $WTA_{1042\text{-}III}$ monomer and for the two selected structures shown in **Fig. 5f** (grey).

**Figure S5. H-bond network and atom distance monitoring for models of CBD500 complexes with $WTA_{1020}$, $WTA_{1042\text{-}I}$ and $WTA_{1042\text{-}III}$ monomers.** Close-up view of selected MD-frames based on better fitting with STD experimental data and closest proximity to MD-cluster centroid structure for $WTA_{1020}$ (presented one of the two from **Fig. 5c**, both from the same cluster) (**a**), $WTA_{1042\text{-}I}$ (presented both of **Fig. 5e**, each from different cluster and with different H-bond pattern) (**b**), and $WTA_{1042\text{-}III}$ (presented one of the two from **Fig. 5f**, both from the same cluster) (**c**). H-bonds are shown as green dashed lines. For perspective clarity, carbon atoms of CBD500 amino acids are colored in green (proximal) or light blue (distant) and those of the ligand in dark grey. Oxygen atoms are colored in red, nitrogen in blue, phosphorus in orange and (polar) hydrogen in light grey. Protein residues and ligand atoms that are involved in H-bonding are labeled in black (CBD500), purple (GlcNAc), yellow (Rbo) or blue (Gal). The right panels correspond to atom distance monitoring in Å for selected H-bonds throughout the 400-ns MD. The dashed lines correspond to the detection limit of 3 Å established to determine the presence of direct H-bonds between CBD500 and the monomeric ligand.

**Figure S6. Comparison of different binding modes of SH3b domain in complex with ligands. a,** Surface representation of the CBD500 in complex with the $WTA_{1042\text{-}III}$ monomer. The proximal and distal domain are colored green and cyan, respectively. **b**, Surface representation of two lysostaphin SH3b domains with the P4-G5 ligand (PDB: 6RK4, Cα atoms 401-493). The SH3b domain and a symmetry-related partner are colored yellow and gray, respectively. **c**, Surface representation of tandem SH3b repeats (colored dark blue and orange) of the CBD of LysPBC5 lysin (PDB ID: 6ILU, Cα atoms 205-339). The PG-interacting residues on both sides of the CBD are highlighted in pink. **d**, Surface representation of tandem SH3b repeats (colored raspberry and limegreen) of the CBD of Psm lysin (PDB ID: 4KRT, Cα atoms 210-342). The structure is rotated ~80 degree to reveal the bound ligand (malonic acid) on the ends of the CBD complex.

Figure S1

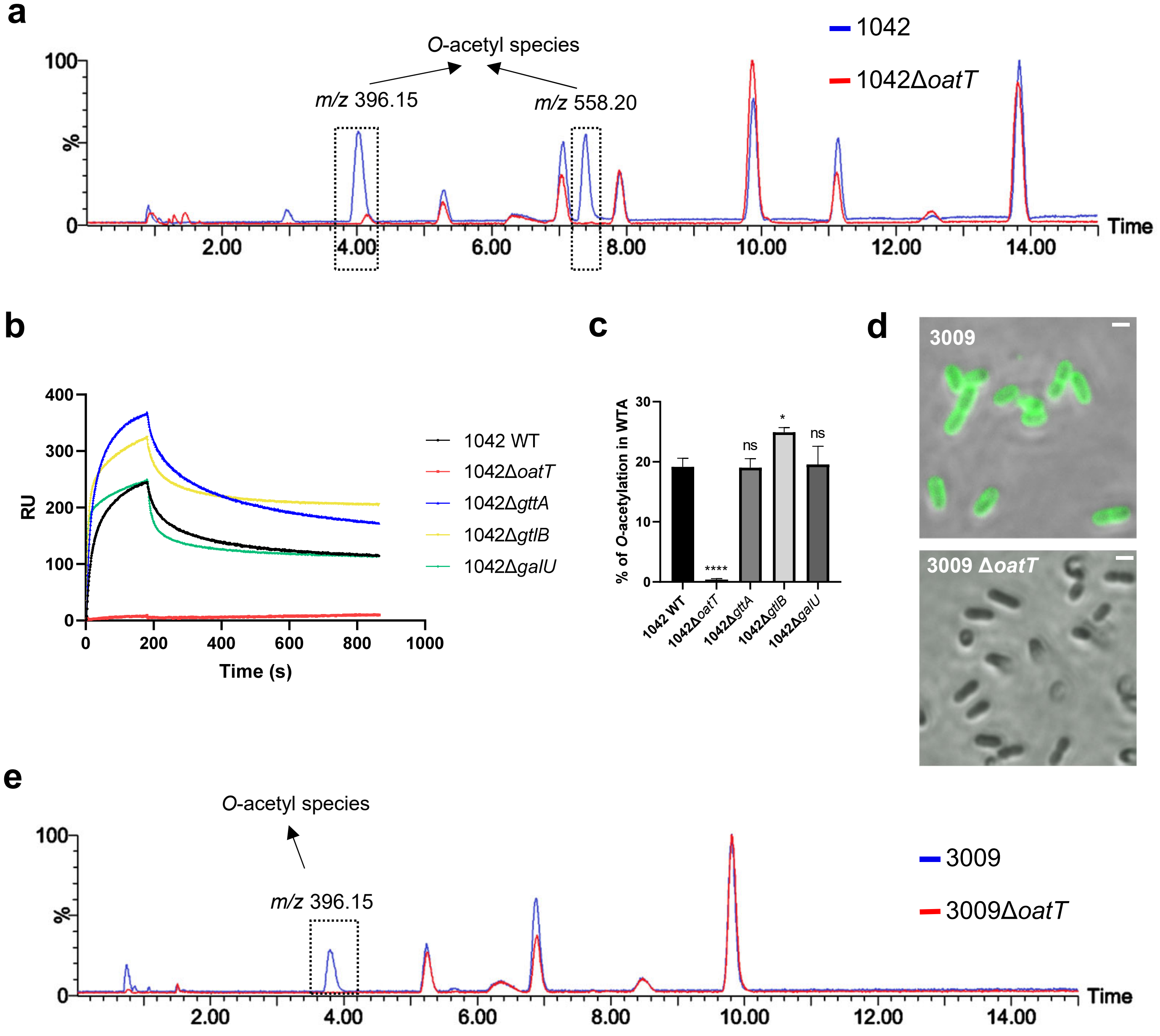
a
O-acetyl species
m/z 396.15
m/z 558.20
1042
1042ΔoatT
100
%
0
2.00
4.00
6.00
8.00
10.00
12.00
14.00
Time
b
400
300
200
100
0
RU
0
200
400
600
800
1000
Time (s)
1042 WT
1042ΔoatT
1042ΔgttA
1042ΔgtlB
1042ΔgalU
c
% of O-acetylation in WTA
30
20
10
0
****
ns
*
ns
1042 WT
1042ΔoatT
1042ΔgttA
1042ΔgtlB
1042ΔgalU
d
3009
3009 ΔoatT
e
O-acetyl species
m/z 396.15
3009
3009ΔoatT
100
%
0
2.00
4.00
6.00
8.00
10.00
12.00
14.00
Time

Figure S2

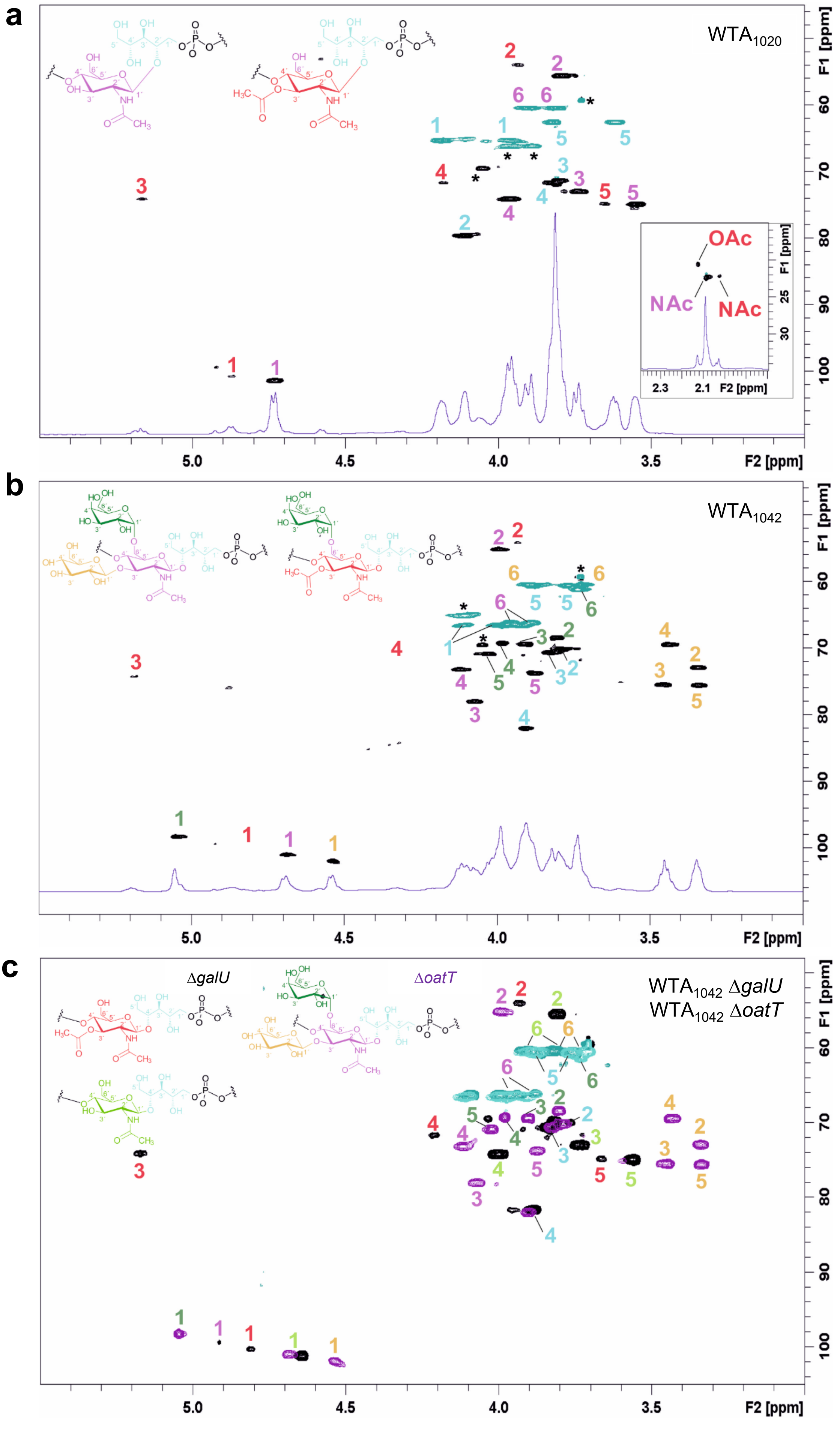

a
WTA1020
OAc
NAc
NAc
2.3
2.1
F2 [ppm]
F1 [ppm]
F2 [ppm]
b
WTA1042
F1 [ppm]
F2 [ppm]
c
ΔgalU
ΔoatT
WTA1042 ΔgalU
WTA1042 ΔoatT
F1 [ppm]
F2 [ppm]

Figure S2

d

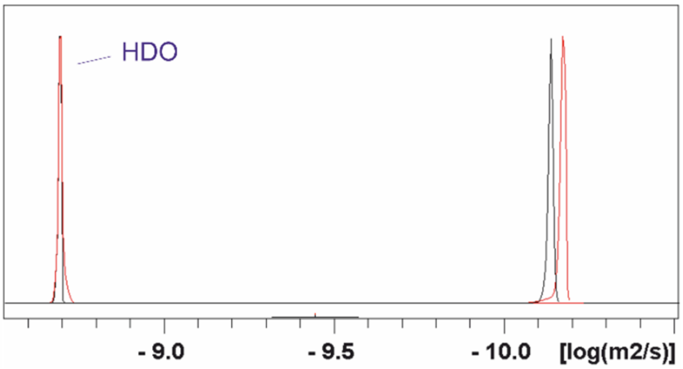
HDO
- 9.0
- 9.5
- 10.0
[log(m2/s)]

e

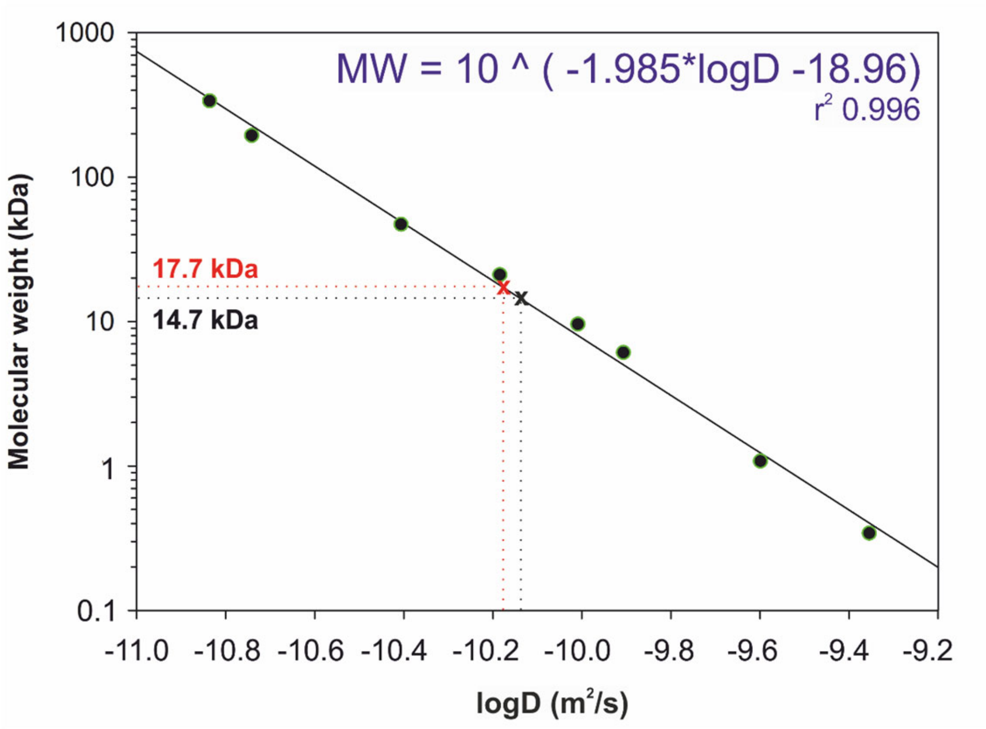
MW = 10 ^ ( -1.985*logD -18.96)
r2 0.996
17.7 kDa
14.7 kDa
1000
100
10
1
0.1
Molecular weight (kDa)
-11.0
-10.8
-10.6
-10.4
-10.2
-10.0
-9.8
-9.6
-9.4
-9.2
logD (m2/s)

Figure S3

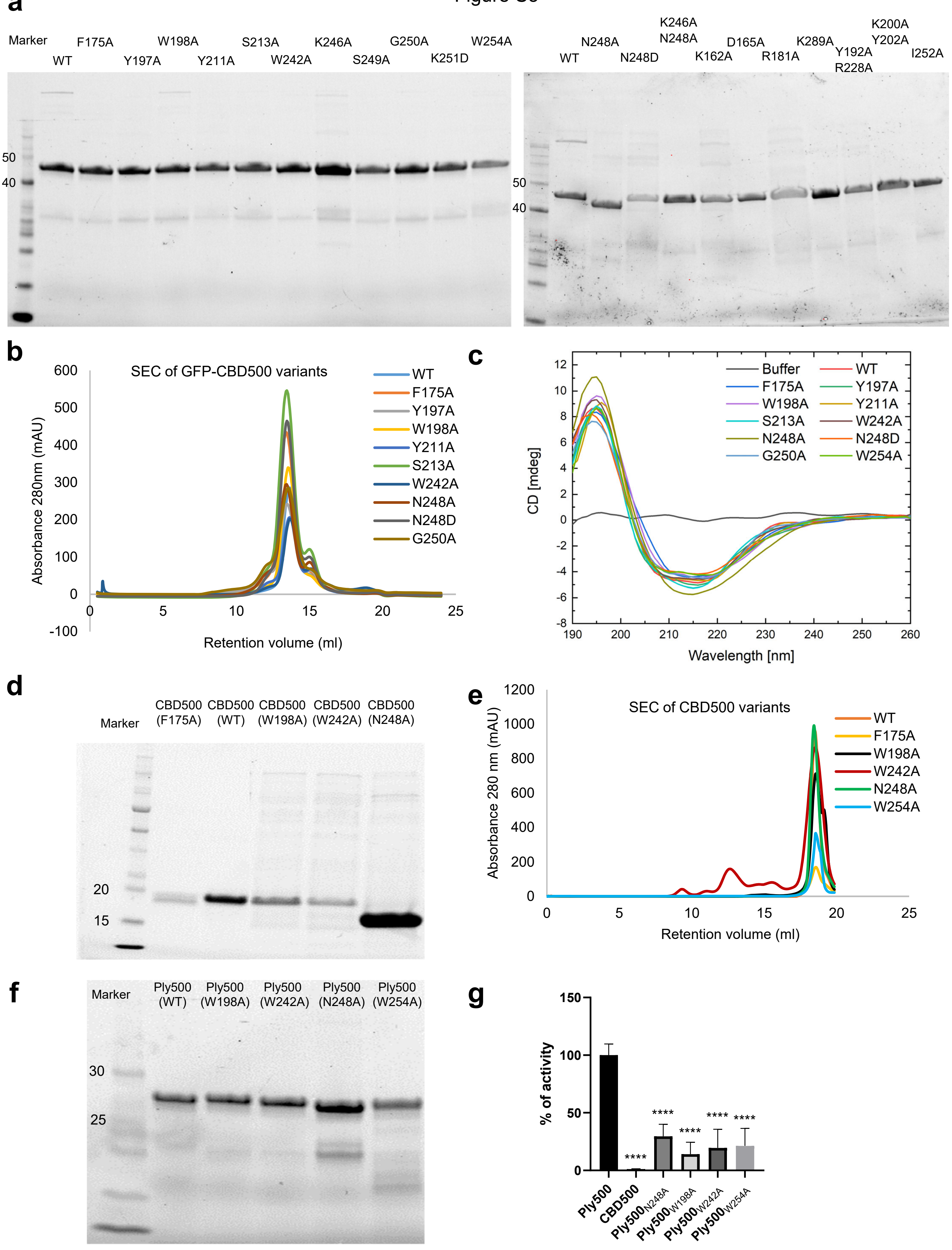
a
Marker
WT
F175A
Y197A
W198A
Y211A
S213A
W242A
K246A
S249A
G250A
K251D
W254A
50
40
WT
N248A
N248D
K246A
N248A
K162A
D165A
R181A
K289A
Y192A
R228A
K200A
Y202A
I252A
50
40
b
SEC of GFP-CBD500 variants
Absorbance 280nm (mAU)
Retention volume (ml)
600
500
400
300
200
100
0
-100
0
5
10
15
20
25
WT
F175A
Y197A
W198A
Y211A
S213A
W242A
N248A
N248D
G250A
c
CD [mdeg]
Wavelength [nm]
12
10
8
6
4
2
0
-2
-4
-6
-8
190
200
210
220
230
240
250
260
Buffer
WT
F175A
Y197A
W198A
Y211A
S213A
W242A
N248A
N248D
G250A
W254A
d
Marker
CBD500 (F175A)
CBD500 (WT)
CBD500 (W198A)
CBD500 (W242A)
CBD500 (N248A)
20
15
e
SEC of CBD500 variants
Absorbance 280 nm (mAU)
Retention volume (ml)
1200
1000
800
600
400
200
0
0
5
10
15
20
25
WT
F175A
W198A
W242A
N248A
W254A
f
Marker
Ply500 (WT)
Ply500 (W198A)
Ply500 (W242A)
Ply500 (N248A)
Ply500 (W254A)
30
25
g
% of activity
150
100
50
0
****
****
****
****
****
Ply500
CBD500
Ply500N248A
Ply500W198A
Ply500W242A
Ply500W254A

Figure S4

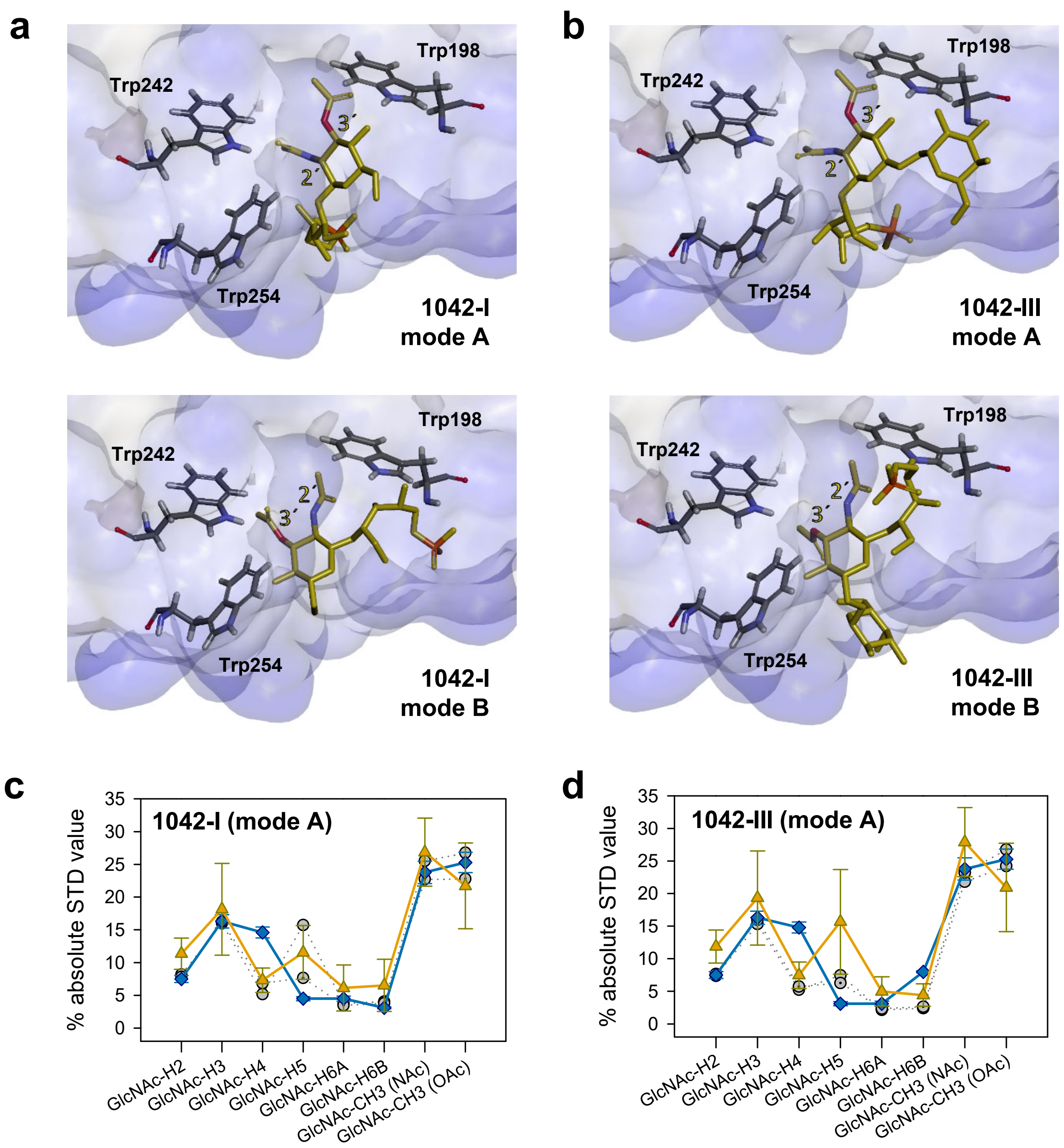
a
Trp242
Trp198
Trp254
1042-I
mode A
Trp242
Trp198
Trp254
1042-I
mode B
b
Trp242
Trp198
Trp254
1042-III
mode A
Trp242
Trp198
Trp254
1042-III
mode B
c
1042-I (mode A)
% absolute STD value
35
30
25
20
15
10
5
0
GlcNAc-H2
GlcNAc-H3
GlcNAc-H4
GlcNAc-H5
GlcNAc-H6A
GlcNAc-H6B
GlcNAc-CH3 (NAc)
GlcNAc-CH3 (OAc)
d
1042-III (mode A)
% absolute STD value
35
30
25
20
15
10
5
0
GlcNAc-H2
GlcNAc-H3
GlcNAc-H4
GlcNAc-H5
GlcNAc-H6A
GlcNAc-H6B
GlcNAc-CH3 (NAc)
GlcNAc-CH3 (OAc)

Figure S5

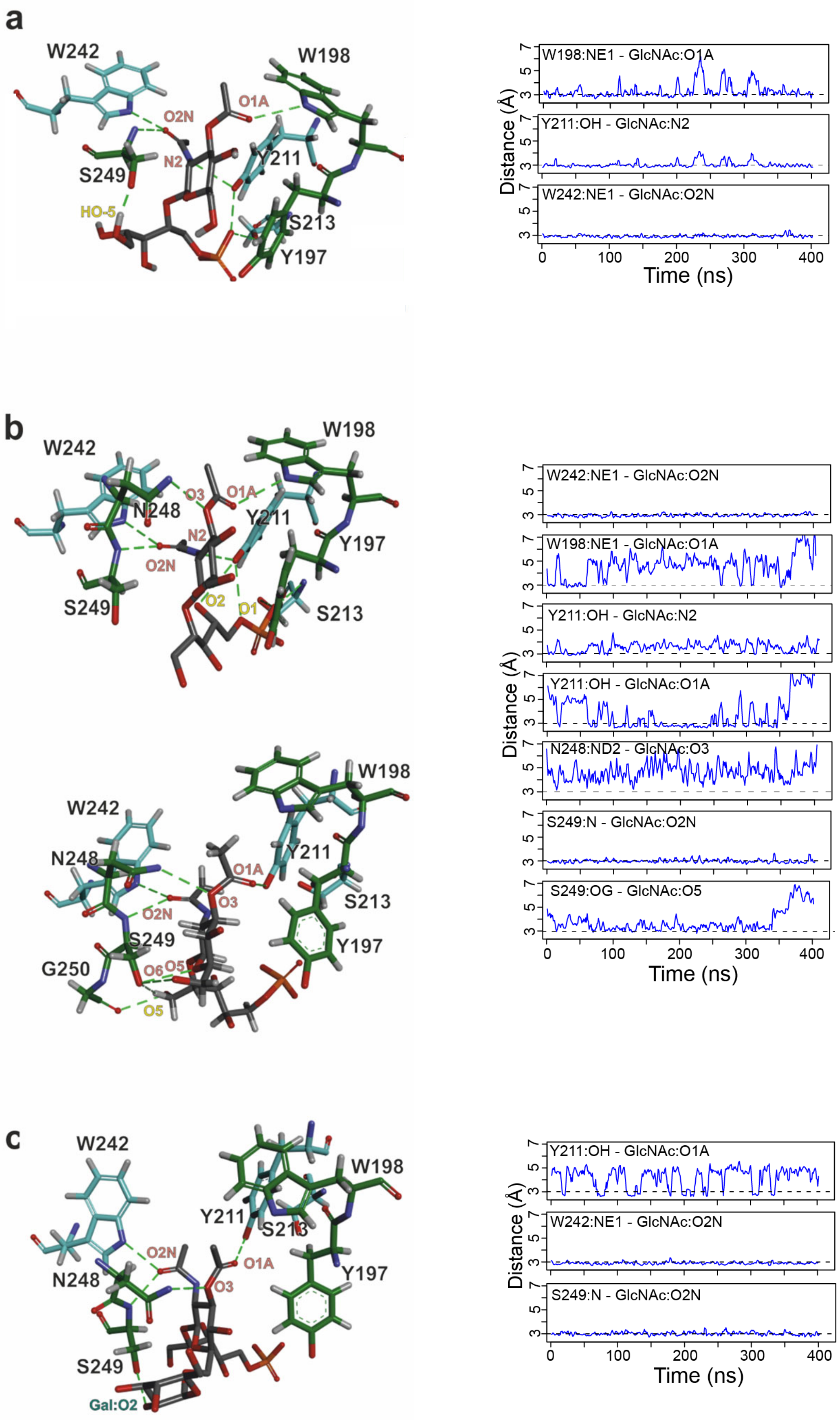
a
W242
W198
O2N
O1A
N2
Y211
S249
HO-5
S213
Y197
W198:NE1 - GlcNAc:O1A
Y211:OH - GlcNAc:N2
W242:NE1 - GlcNAc:O2N
Distance (Å)
Time (ns)
b
W242
W198
N248
O3
O1A
Y211
N2
Y197
O2N
S249
O2
O1
S213
G250
O6
O5
W242:NE1 - GlcNAc:O2N
W198:NE1 - GlcNAc:O1A
Y211:OH - GlcNAc:N2
Y211:OH - GlcNAc:O1A
N248:ND2 - GlcNAc:O3
S249:N - GlcNAc:O2N
S249:OG - GlcNAc:O5
Distance (Å)
Time (ns)
c
W242
W198
Y211
S213
O2N
O1A
N248
O3
Y197
S249
Gal:O2
Y211:OH - GlcNAc:O1A
W242:NE1 - GlcNAc:O2N
S249:N - GlcNAc:O2N
Distance (Å)
Time (ns)

Figure S6

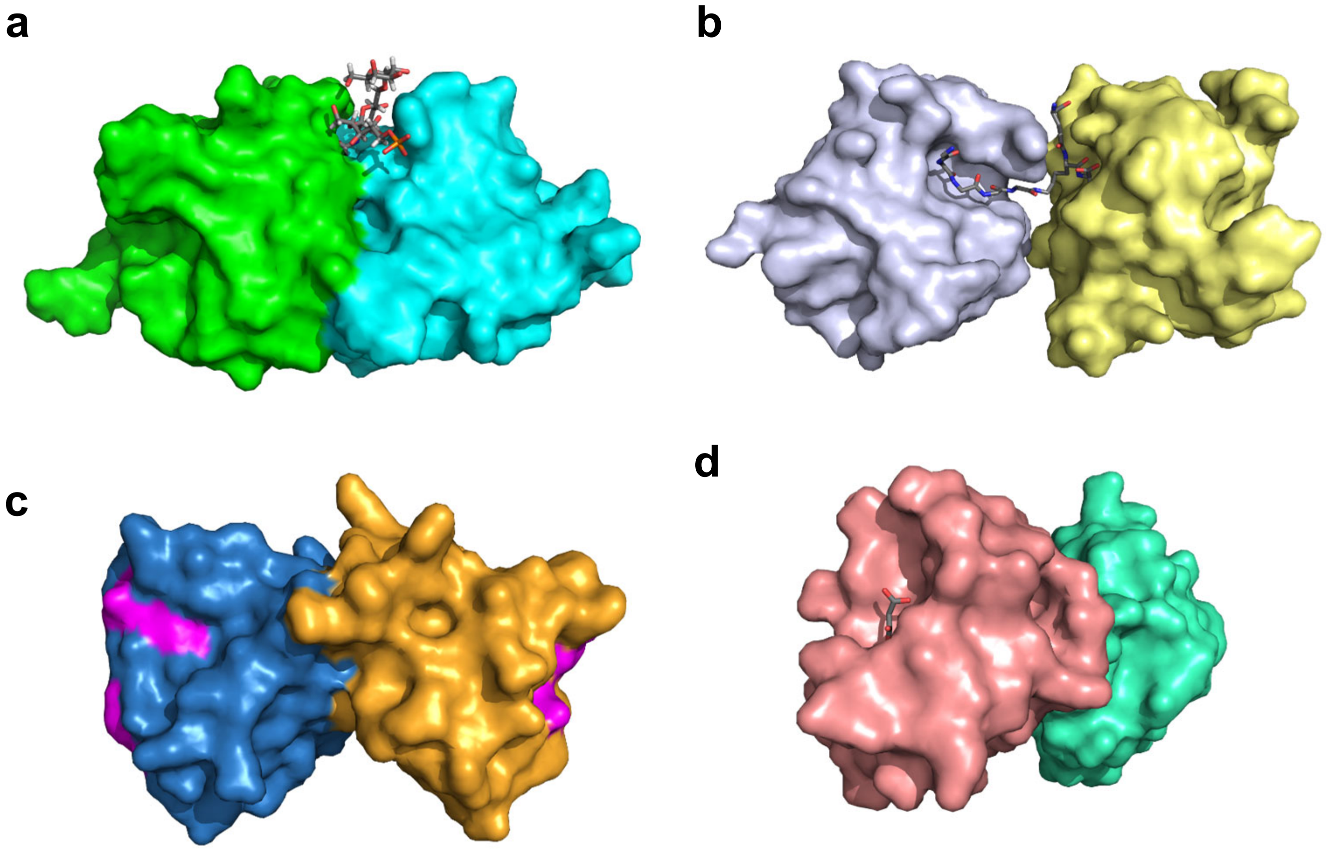
